# Burden of tumor mutations, neoepitopes, and other variants are dubious predictors of cancer immunotherapy response and overall survival

**DOI:** 10.1101/665026

**Authors:** Mary A. Wood, Benjamin R. Weeder, Julianne K. David, Abhinav Nellore, Reid F. Thompson

**Affiliations:** Computational Biology Program, Oregon Health & Science University; Portland VA Research Foundation; Department of Biomedical Engineering, Oregon Health & Science University; Department of Surgery, Oregon Health & Science University; Department of Radiation Medicine, Oregon Health & Science University; Department of Medical Informatics & Clinical Epidemiology, Oregon Health & Science University; Division of Hospital and Specialty Medicine, VA Portland Healthcare System

**Keywords:** tumor mutational burden, TMB, neoepitopes, neoepitope burden, neoantigens, splice junctions, retained introns, tumor variant burden, immunotherapy response

## Abstract

**Background:** Tumor mutational burden (TMB, the quantity of aberrant nucleotide sequences a given tumor may harbor) has been associated with response to immune checkpoint inhibitor therapy and is gaining broad acceptance as a result. However, TMB harbors intrinsic variability across cancer types, and its assessment and interpretation are poorly standardized.

**Methods:** Using a standardized approach, we quantify the robustness of TMB as a metric and its potential as a predictor of immunotherapy response and survival among a diverse cohort of cancer patients. We also explore the additive predictive potential of RNA-derived variants and neoepitope burden, incorporating several novel metrics of immunogenic potential.

**Results:** We find that TMB is a partial predictor of immunotherapy response in melanoma and non-small cell lung cancer, but not renal cell carcinoma. We find that TMB is predictive of overall survival in melanoma patients receiving immunotherapy, but not in an immunotherapy-naive population. We also find that it is an unstable metric with potentially problematic repercussions for clinical cohort classification. We finally note minimal additional predictive benefit to assessing neoepitope burden or its bulk derivatives, including RNA-derived sources of neoepitopes.

**Conclusions:** We find sufficient cause to suggest that the predictive clinical value of TMB should not be overstated or oversimplified. While it is readily quantified, TMB is at best a limited surrogate biomarker of immunotherapy response. The data do not support isolated use of TMB in renal cell carcinoma.

## BACKGROUND

The advent of immunotherapy as a promising form of cancer treatment has been accompanied by a parallel effort to explore potential mechanisms and drivers of therapeutic response. For instance, tumor mutational burden (TMB, the overall quantity of aberrant nucleotide sequences a given tumor may harbor) has been associated with response to immune checkpoint inhibitor therapy (1) and overall survival (2). Similarly, the quantity of non-synonymous single nucleotide variants was shown to be associated with immunotherapy response in several independent clinical cohorts (3–6). Other sources of sequence variation such as frameshifting insertions/deletions (7) and tumor-specific alternative splicing (e.g. intron retention (8)) have also been found to correlate with immunotherapy response. These phenomena are widely accepted and appear to be particularly pronounced in patients harboring DNA repair deficiencies (9). Indeed, the checkpoint inhibitor, pembrolizumab, was granted accelerated disease-agnostic approval by the FDA on this basis for any cancer patient harboring deficiencies in their capacity to perform DNA mismatch repair (10). Moreover, an expanding cohort of clinical immunotherapy trials (e.g. NCT03668119, NCT03178552, NCT03519412) are actively utilizing TMB status as a key inclusion criterion. However, there is wide variability among techniques for measuring and interpreting TMB, raising questions of utility and reproducibility (11).

Given the perceived critical importance of TMB in the research setting and its emerging role in the oncology clinic, we sought to quantify the robustness of TMB as a metric, and explore its deeper nuances using pooled whole exome sequencing data from a variety of previously published studies. While TMB is generally correlated with downstream metrics such as neoepitope burden, we also explore the predictive capacity of neoepitope burden and its derivatives including adjustment for MHC binding robustness and peptide sequence novelty, as well as RNA-derived sources of neoepitopes.

## METHODS

### Variant Identification and Neoepitope Prediction

We assembled a cohort of 457 tumor samples from 431 different cancer patients from publicly available data, including 302 melanoma patients (326 tumor samples) (1,4–6,12–16), 34 non-small cell lung cancer (NSCLC) patients (34 tumor samples) (3), 10 prostate cancer patients (10 tumor samples) (17), 57 renal cell carcinoma (RCC) patients (58 tumor samples) (18), and 28 mismatch repair (MMR) deficient (as determined by polymerase chain reaction or immunohistochemistry (9)) colon, endometrial, and thyroid cancer patients (29 tumor samples) (9) (see Supplementary Table 1). Despite attempts to obtain these data, we unfortunately were forced to omit tumor samples from 75 NSCLC patients (19), for whom data was not available due to limitations of patient consent at the time of the study. Alignment of whole exome sequencing (WES) reads was performed as described previously (23).The Mbp of genome covered was determined using bedtools genomecov (v2.26.0) (24), where any base covered by a depth of at least 6 reads was considered covered, as this is twice the minimum read depth required for variant detection by SomaticSniper (25) and VarScan 2 (26). Somatic and germline variant calling were performed as described previously (23). To obtain coverage-adjusted mutation burdens for each patient, we divided the number of consensus somatic variants by the Mbp of genome covered by sequencing. We employed HapCUT2 for patient-specific haplotype phasing. To do this, germline and consensus somatic variants were combined into a single VCF using `neoepiscope`’s (23) (v0.3.5) merge functionality. HapCUT2’s extractHAIRS software was run with the merged VCF and the tumor alignment file, allowing for extraction of reads spanning indels, to produce the fragment file used with HapCUT2 to predict haplotypes. Neoepitopes of 8-24 amino acids in length were predicted for this cohort using `neoepiscope`, including background germline variation and variant phasing, and enumerating neoepitopes from protein coding, nonsense mediated decay, polymorphic pseudogene, T cell receptor variable, and immunoglobulin variable transcripts. Additionally, to better understand how the choice of variant caller impacts downstream neoepitope predictions, we ran `neoepiscope` excluding background germline variation and variant phasing separately for our consensus somatic variants and variants produced by individual variant calling tools, only enumerating neoepitopes from protein coding transcripts. For patients with multiple tumor samples, the median mutation and neoepitope burdens across samples were retained. Variants that were pathogenic or likely pathogenic in cancer according to ClinVar (34) were identified using Open-CRAVAT (35), and neoepitopes deriving from these variants were flagged. We used the software mSINGS (36) (bit bucket commit 030289381f3b7aee24d8eccbb69b3e66711f5bb0) to identify tumors with MSI positive status. The software was run on each tumor alignment file, and the provided TCGA msi_bed, msi_baseline, msi_intervals were used.

### RNA Variant Identification

Among the overall cohort, 106 patients (89 melanoma patients (1,4–6) and 17 RCC patients (18)) had complementary tumor RNA-sequencing (RNA-seq) data. We aligned RNA-seq reads to both the GRCh37d5 and GRCh38 genomes using STAR (v2.6.1c) (37), using the ‘intronMotif’ --outSAMstrandField option and specifying NH, HI, AS, nM, and MD fields with the --outSAMattributes option. To identify putative tumor-specific splice junctions, we first downloaded called junction data including coverage and bed files for TCGA and GTEx using recount2 (38). GENCODE version 28 annotations (39) were downloaded and parsed to collect full coordinates and left and right splice sites of junctions from annotated transcripts. The TCGA phenotype file from Rail-RNA (40) was parsed to collect sample type (primary, recurrent, or metastatic tumor vs. matched normal). A new SQLite3 database was created to index all GTEx and TCGA junctions, with linked tables containing 1) sample ids and associated junction ids; 2) sample ids and phenotype information for each sample; and 3) junction ids and junction information including GENCODE annotation status and location within protein coding gene boundaries. Junctions were extracted from the SJ.out output files generated by STAR; only junctions with canonical splice motifs (GT-AG, GC-AG, and AT-AC) were collected. No minimum read count was imposed for a junction to be called in a sample. The known junction index was queried to collect all junctions found in normal tissue either in GTEx or in TCGA matched normal samples and these normal junctions were filtered out from the single sample set. We used the MetaSRA (41) web query interface to collect Sequence Read Archive (SRA) accession numbers for non-cancerous melanocyte cell line (42) and primary cell (43) RNA-seq experiments. The resulting accession numbers were queried against the Snaptron junction database (44,45) to download junctions from across the entire genome. All junctions found in these normal melanocyte samples as well as all fully GENCODE-annotated junctions were also eliminated from each single-sample junction set. Again, no minimum read support was required; a single read covering a junction in a single non-cancer sample (SRA, GTEx, or TCGA) eliminated the junction from the patient set. Finally, we removed junctions where neither end was found in GENCODE-annotation, yielding a list of putative tumor-specific splice sites for each patient. Supplementary Figure 1 (rows 2-4) illustrates the variety of splicing alterations captured.

We identified tumor-specific retained introns (see Supplementary Figure 1, row 5) using Keep Me Around (kma) (46). We aligned RNA-seq reads to a modified version of the GRCh37d5 using Bowtie 2 (v2.3.4.3) (47), and quantified reads using eXpress (v1.5.1) (48), as per kma recommendations. After computing intron retention, we used kma’s filters to retain only transcripts that were expressed at greater than or equal to 1 transcripts per million (TPM) in at least 25% of samples, transcripts that had at least 5 unique counts in at least 25% of samples, and transcripts that had greater than 0 and less than 100 percent of introns retained. To prevent inclusion of artifacts from unprocessed transcripts, we identified outlier introns among the distribution of transcript read counts, only retaining introns with a read count greater than 3 median absolute deviations above the median intron read count for a transcript, and greater than or equal to the read count for the transcript itself. To filter out retained introns that may be expressed in normal tissues, we performed the same analysis using using publically available RNA-seq reads from melanocyte samples of 106 newborns (49). Any retained introns identified from the melanocyte RNA-seq data were then removed from the retained introns identified from the tumor RNA-seq data. Neoepitopes deriving from retained introns were predicted using the reading frame from the 5’ end of the transcript of origin prior to the intron, enumerating peptides 8-24 amino acids in length.

### HLA Type Prediction and Related Analyses

MHC Class I alleles for each patient were predicted from tumor WES reads using Optitype (v1.0) (50), and MHC Class II alleles for each patient were predicted from tumor WES reads using seq2hla (v2.2) (51). For each neoepitope sequence predicted from phased variants (see above), a patient’s predicted MHC Class I and MHC Class II alleles were used for binding affinity predictions with MHCnuggets (v2.1) (52). Neoepitopes were counted toward a patient’s neoepitope burden if they bound at least one of a patient’s MHC alleles with high affinity (<= 500 nM). For comparison with neoepitope burdens reported by the authors of the five original manuscripts with reported neoepitope burdens, we tallied binding predictions separately based on their methodology, using binding affinity predictions from NetMHCpan (v4.0) (53) for a more direct comparison. For patients from the studies by Carreno et al. (4) and Rizvi et al. (3) we considered only 9mer epitopes; for patients from the study by Van Allen et al. (1) we considered only 9mer and 10mer epitopes; and for the studies by Hugo et al. (5) and Roh et al. (14) we considered 9mer, 10mer, and 11mer epitopes. For the epitopes from patients from the Carreno et al. study, we only considered binding to HLA-A*02:01 as in their paper, while for the other studies we considered binding to any MHC Class I epitope. Additionally, we determined the burden of processed neoepitopes (those predicted to be cleaved by the proteasome, transported by TAP, and presented on the cell surface by an MHC Class I molecule) using NetCTLpan (v1.1) (54). For each tumor sample, we ran NetCTLpan predictions for all 8mer, 9mer, 10mer, and 11mer neopeptides with each MHC Class I epitope predicted by Optitype. A neopeptide was counted toward the burden of processed epitopes if its NetCTLpan combined score rank was in the top 1% for at least one MHC allele.

### Modified Neoepitope Burden

To better understand how different features of tumor neoepitopes might influence response to immunotherapy, we produced several normalized neoepitope burdens. We first calculated neoepitope burden for each patient weighted by MHC allele presentation, where a predicted neoepitope sequence counted toward the patient’s neoepitope burden once for each of the patient’s MHC alleles that was predicted to bind that neoepitope with high affinity (<= 500 nM). Second, neoepitope burden was calculated for each patient weighted by amino acid mismatch as follows. The closest normal peptide in the human proteome to each neoepitope was identified using blastp (v2.6.0) (55), selecting for lowest E value or, in the case of a tie among multiple peptide sequences, the selected peptide was that with the highest weighted BLOSUM62 similarity (as described previously (56)). A neoepitope sequence was counted toward the patient’s neoepitope burden once for each amino acid mismatch between the neoepitope and its closest normal peptide. Third, neoepitope burden was calculated for each patient weighted by TCGA transcript expression of the transcript(s) of origin for each neoepitope. We identified expressed transcripts in matched TCGA cancer types for each disease type in our cohort (SKCM for melanoma, LUAD/LUSC for NSCLC, COAD for colon cancer, UCEC for endometrial cancer, THCA for thyroid cancer, PRAD for prostate cancer, and KIRC for RCC) from TPM values generated by the National Cancer Institute (57). A transcript was considered “expressed” for a cancer type if the 75th quantile TPM value for that transcript in that disease was greater than 1 TPM. Because these TPM values were based on GRCh38 transcripts, we used liftOver (58) to convert the coordinates of a neoepitope’s mutation of origin to GRCh38 coordinates and identify overlapping transcripts. A neoepitope sequence was counted toward the patient’s neoepitope burden once for each transcript of origin expressed in TCGA. Note that for patients with tumor RNA-seq data (see above), we also calculated neoepitope burden weighted by patient-specific expression of the transcript(s) of origin for each neoepitope. We used Rail-RNA (v0.2.4b) (40) on RNA-seq alignments to the GRCh37d5 genome to identify covered exons, and a transcript was considered “expressed” if at least 1 read covered any exon in the transcript. A neoepitope sequence was counted toward the patient’s neoepitope burden once for each expressed transcript of origin. Finally, we multiplicatively combined these weighted burdens by multiplying scores for each epitope and totaling all epitope scores: allele presentation score by amino acid mismatch score, allele presentation score by TCGA expression score, allele presentation score by patient-specific expression score (if relevant), amino acid mismatch score by TCGA expression score, amino acid mismatch score by patient-specific expression score (if relevant), allele presentation score by amino acid mismatch score by TCGA expression score, and allele presentation score by amino acid mismatch score by patient-specific expression score (if relevant).

### Statistical Analysis

Statistical analysis was performed in R (v3.5.1). The rlm function from the MASS package (v7.3-51.4) was used for robust linear model fitting, and the cor.test function was used for determining Pearson product-moment correlation values. To determine variability in TMB across variant calling tools, the median of pairwise differences in TMB between tools was divided by the median TMB across tools for each patient; the median of these values across patients was reported. The roc function from the pROC package (v1.14.0) was used to generate ROC curves for any predictors of immunotherapy response and to determine their AUC for all patients with reported immunotherapy response status (409/414, after excluding 3 colon cancer, 1 prostate cancer, and 1 RCC patient that lacked documented response status). Logistic regression was performed using the glm function to model therapeutic response as a linear function of TMB (on log scale), and neoepitopes (log scale) on the 245 melanoma patients, 50 RCC patients, and 33 NSCLC patients with reported immunotherapy response status to either aCTLA4 or aPD1 treatment alone (excluding dual/combination checkpoint inhibitor therapy). For the subset of these patients with available RNA-seq data (see Supplementary Table 1), tumor variant burden (TVB; the sum of somatic variants, tumor-specific splice junctions, and tumor-specific retained introns; log2 scale) was also modeled. The fit models were subsequently used to estimate the odds of therapeutic response at the 25^th^ and 75^th^ TMB, TVB, and neoepitope percentiles. Each cancer type was modeled separately, with the melanoma model accounting for differences in aCTLA4 vs. aPD1 response rates. P-values were adjusted for multiple comparisons using the Benjamini-Hochberg method with the p.adjust function.

### Survival Analysis

Due to the low number of observed events for some cancers, only melanoma and RCC patient cohorts were appropriate for survival analysis. Patients were included in survival analysis if they had information on both overall survival status, as well as either time to event or time to censorship data. In total, 218 melanoma patients and 56 RCC patients were selected for analysis in R (v3.5.1). The coxph function from the survival package (v2.44-1.1) was used to fit proportional hazards regression models, and the survfit function from the survival package was used to compute survival curves. For comparison with patients not treated with immunotherapy, we also performed survival analysis with SKCM and KIRC patients from TCGA. We obtained mutation annotation format (MAF) files and clinical data for these patients from the Broad Institute (59). Patients with both mutation information and survival information were used for analysis (320 SKCM patients and 415 KIRC patients). Mutational burden was determined by counting the number of somatic variants listed in each patient’s MAF file, and a patient was considered to have survival information if they had information on time to death or a non-zero and non-negative value on time to last follow up.

## RESULTS

### Distribution of tumor variant and neoepitope burdens

We find that the median TMB (based on consensus DNA variant calls; see Methods) varies by an order of magnitude across disease types, ranging from 635.5 variants for prostate cancer to 5,632.5 variants for MMR-deficient cancers (Supplementary Figure 2). Adjusting by genome coverage for each patient (see Methods), the median TMB was 18.03 mutations/Mbp of genomic coverage (ranging from 5.17 for RCC prostate cancer to 26.82 for MMR-deficient cancers, see Figure 1A). The majority of variants were found to be single nucleotide variants (median 85.07% per patient), with the remainder from in-frame and frameshift insertions and deletions (ranging from a median 7.52% indels for RCC to 37.26% indels for prostate cancer, see Supplementary Figure 3). Note that RNA variants such as alternative exon-exon junctions and retained introns were also assessed in the subset of patients with corresponding RNA-sequencing data (see Methods). Overall, tumor-specific junction burden appeared to be less variable across cancer types (ranging from 1301 for RCC to 2048.5 for melanoma). While retained introns (RI) have also been described as a source of neoepitopes (8), only 27 melanoma patients with RNA-seq data had any predicted RIs, with a median RI burden in those patients of 929 introns. Integrating these tumor DNA and RNA variants (given matched RNA-seq data) into a single combined tumor variant burden (TVB) yielded a median increase of 2345 variants per patient, with RNA sources of variation accounting for an average 40.8% of overall variants (see Figure 2). Moreover, consideration of DNA variant burden alone neglects substantial somatic variation for some patients, as RNA sources of variation can constitute up to 86.7% of TVB.

**Figure 1:**
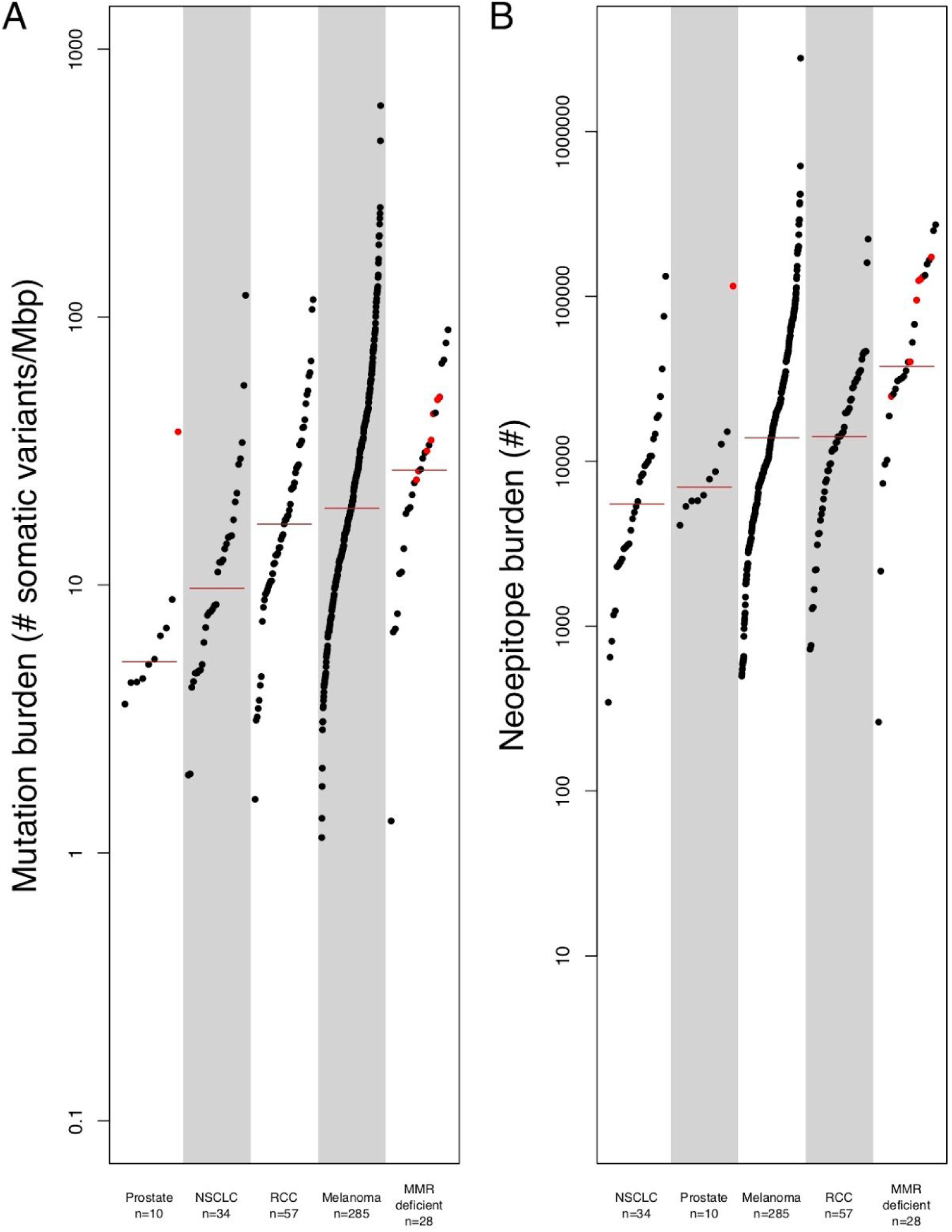
Per-patient distribution of mutation and neoepitope burdens across 7 cancer types. A) The number of somatic DNA variants per patient (scaled for sequence coverage) are shown along the y-axis, with each dot representing an individual cancer patient (cancer types shown along the x-axis). Note that MMR-deficient cancers here represent a cohort of 3 different cancer types including colon, endometrial, and thyroid with evidence of mismatch repair deficiency as determined by polymerase chain reaction or immunohistochemistry (9). Red colored dots correspond to patients with microsatellite instability as determined by mSINGS (see Methods). B) The number of putative neoepitopes per patient are shown along the y-axis, with each dot representing an individual cancer patient (cancer types shown along the x-axis). Abbreviations as follows: MMR=mismatch repair.

**Figure 2:**
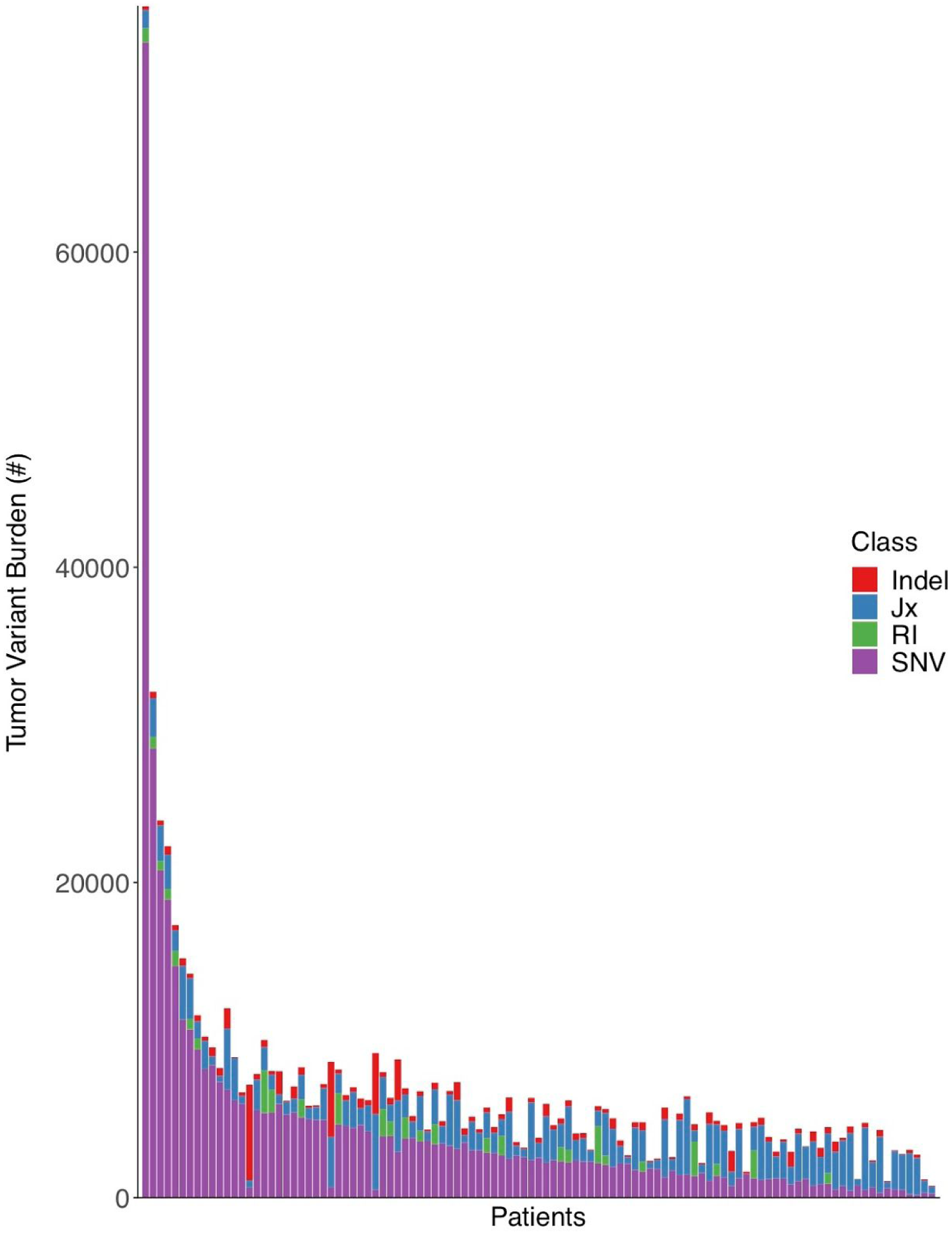
Per-patient distribution of overall tumor variant burden and its components. The number of total tumor variants per patient is shown along the y-axis, with the numbers of retained introns (RI), tumor-specific exon-exon junctions (Jx), insertions/deletions (Indel), and single nucleotide variants (SNV) shown in green, blue, red, and purple, respectively. The data for each individual patient is displayed as stacked bars along the x-axis, sorted from left to right by the number of single nucleotide variants (from highest to lowest).

As TMB and TVB are indirect assessments of cancer neoantigen load, we next calculated DNA-derived, RNA-derived and overall neoepitope burdens per patient from putative protein-level variation (see Methods). The median per-patient DNA-derived neoepitope burden (for peptides predicted to bind to at least one of a patient’s MHC Class I or II alleles) was 13,512 peptides (ranging from 5,511.5 for NSCLC to 37,710.5 for MMR-deficient cancers, see Figure 1B) and was highly correlated with TMB itself (Pearson’s product-moment correlation of 0.63, p < 2.2×10^−16^; see Supplementary Figure 4). There were generally more MHC Class I epitopes than Class II epitopes, with a median Class I epitope burden of 6,337 peptides (ranging from 2,366 for NSCLC to 15,645.5 for MMR-deficient cancers) and a median Class II epitope burden of 6,027 peptides (ranging from 2,167.5 for NSCLC to 23,554.5 for MMR-deficient cancers). We also assessed the burden of Class I epitopes predicted to be processed via the proteasome, transported through TAP, and presented on the cell surface (see Methods), with a median 766 such epitopes per patient (ranging from 366 for prostate cancer and 2,536.5 for MMR-deficient cancers). While not all patients possessed RIs, the median per-patient RNA-derived neoepitope burden among the 27 melanoma patients with predicted RIs (366,843 peptides) was an order of magnitude higher than DNA-derived neoepitopes in the vast majority of cases (Supplementary Figure 5).

In addition to reporting the bulk number of neoepitopes per patient, we also analyzed the distribution of peptide presentation by patient-specific HLA types. Overall, a median of 8.91% of possible peptides are presented by one or more patient-specific MHC Class I or II alleles. Among these, any given neoepitope is, on average, only presented by a single MHC allele (Figure 3A, Supplementary Figure 6A). There are many additional degrees of freedom to surveil the peptide-level consequences of an individual variant (e.g. individual single nucleotide variants may give rise to as many as 272 different peptides of 8-24aa lengths, any of which might be presented via one or more MHC Class I or II alleles). As such, we find that 83.4% of all DNA variants resulting in peptide-level change(s) have at least one neoepitope putatively presented by at least one HLA allele, with a median of 3 different HLA alleles able to present one or more neoepitopes from each individual variant (Figure 3B, Supplementary Figure 6B). Moreover, the percentage of variants presented increases with increasing MHC heterozygosity (Figure 3C, Supplementary Figure 7). Within the cohort, 329 patients had pathogenic cancer-related mutations (see Methods), with an average of 2.8 such variants per patient among those patients. Consistent with prior work demonstrating a relative paucity of peptide presentation from cancer driver mutations (60), we find that a smaller number (approximately 68.5%) of driver variants in this cohort yielded neoepitopes, with only 10.4% of neopeptides from these variants on average being predicted to bind to any of a patient’s HLA alleles (Figure 3).

**Figure 3:**
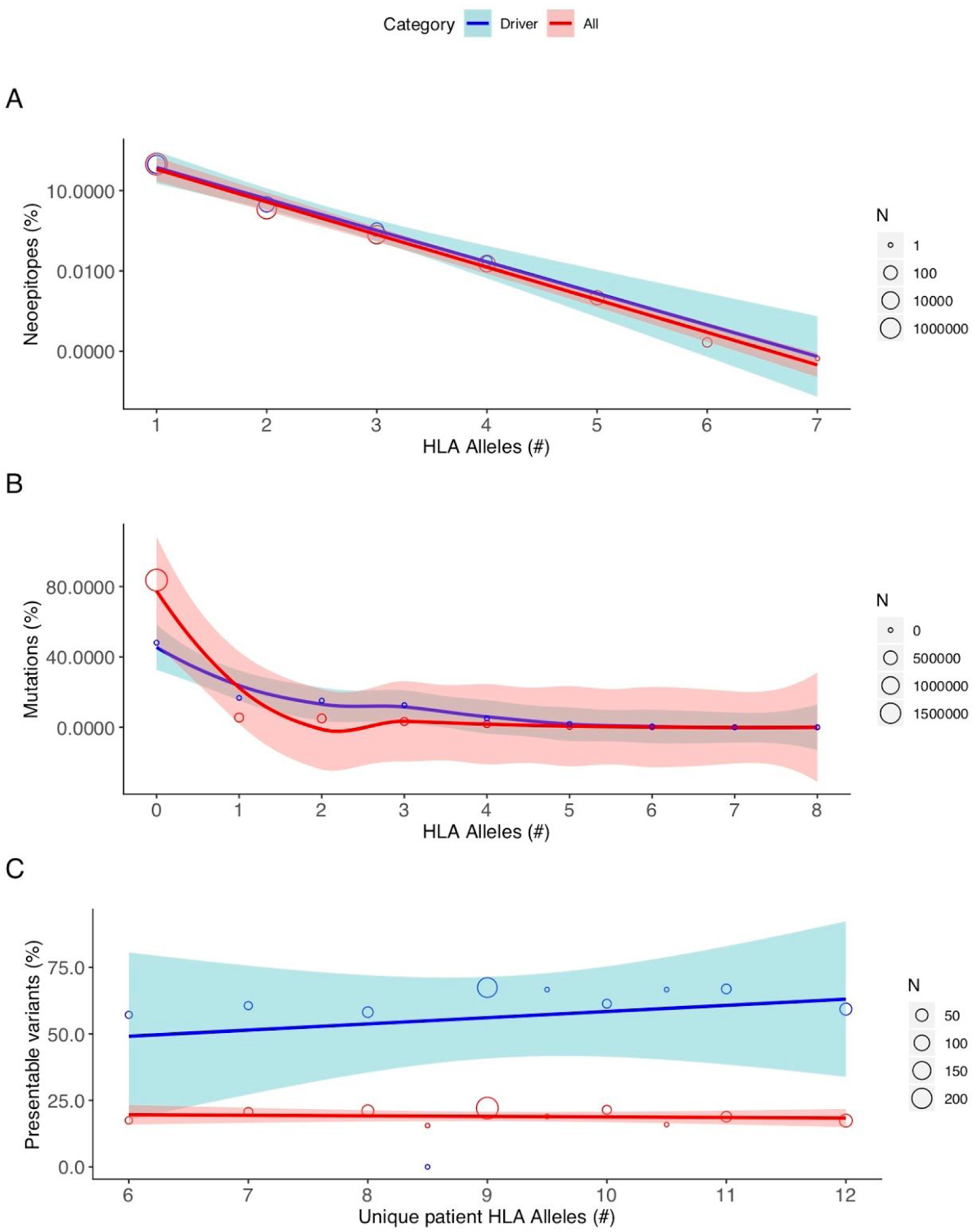
Robustness of putative neoepitope presentation. A) The number of unique patient-matched HLA alleles that are predicted to present an individual neoepitope is shown along the x-axis, with the y-axis (log-scale) corresponding to the overall percent of neoepitopes sharing that same robustness of HLA presentation. Red and blue curves denote the best fit line based on linear regression for all neoepitopes and those resulting from cancer driver mutations, respectively. The surrounding red and light blue shading denotes the 95% confidence interval for all and driver-derived neoepitopes, respectively. Individual data points are shown as open circles, whose diameter corresponds to the number of neoepitopes as shown by the corresponding scale at right. B) The total number of unique patient-matched HLA alleles that are predicted to present one or more neoepitopes arising from a single DNA mutation is shown along the x-axis, with the y-axis corresponding to the overall percent of mutations sharing that same robustness of HLA presentation. Red and blue curves denote the best fit line based on local polynomial regression for all mutations and cancer driver mutations, respectively. The surrounding red and light blue shading denotes the 95% confidence interval for all and driver mutations, respectively. Individual data points are shown as open circles, whose diameter corresponds to the number of mutations as shown by the corresponding scale at right. C) The percentage of total variants that are predicted to be presented by one or more patient-matched HLA alleles is shown along the y-axis, with the x-axis corresponding to the number of unique HLA alleles for that patient. Red and blue curves denote the best fit line based on linear regression for all mutations and cancer driver mutations, respectively. The surrounding red and light blue shading denotes the 95% confidence interval for all and driver mutations, respectively. Individual data points are shown as open circles, whose diameter corresponds to the number of mutations as shown by the corresponding scale at right. Note that a predicted HLA binding affinity threshold of ≤500nM was used in all cases (see Methods).

### Tumor variant and neoepitope burdens as predictors of response and survival

We next sought to quantify immunotherapy response rate as a function of TMB, TVB, and neoepitope burden. Using disease-specific logistic regression models, we found that neither TVB nor neoepitope burden were significant predictors of immunotherapy response (see Table 1). In contrast, for NSCLC patients there was a 120.8% increase in the odds of response per log2 fold change in TMB (p = 0.034), though these results were not significant upon adjustment for multiple hypothesis testing (p=0.053). Similar effects were not seen for melanoma (p = 0.267) or RCC (p = 0.973).

**Table 1:**
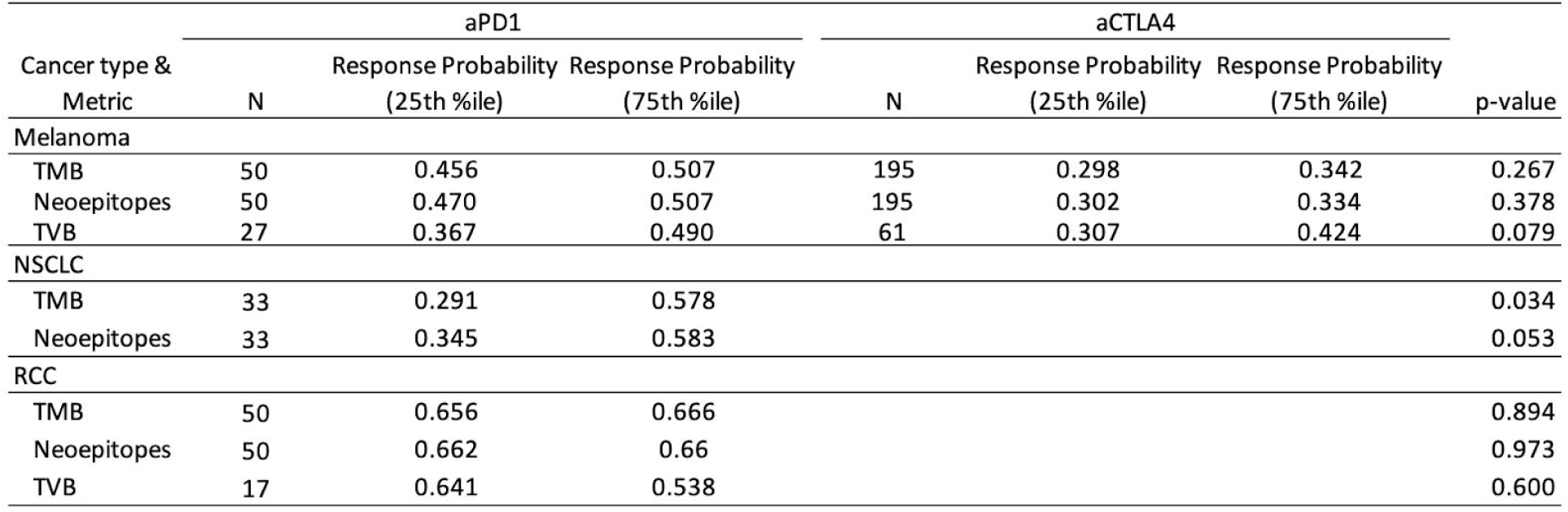
Immunotherapy (αPD1 and αCTLA4) response probability based on logistic regression models of tumor mutational burden (TMB), neoepitope burden (Neoepitopes), and combined tumor DNA- and RNA-variant burden (TVB) for melanoma, non-small cell lung cancer (NSCLC) and renal cell carcinoma (RCC). P-values are reported on a per-model basis without correction for multiple comparisons per cancer type.

| Cancer type & Metric | aPD1 |  |  | aCTLA4 |  |  | p-value |
| --- | --- | --- | --- | --- | --- | --- | --- |
|  | N | Response Probability (25th %ile) | Response Probability (75th %ile) | N | Response Probability (25th %ile) | Response Probability (75th %ile) |  |
| Melanoma |  |  |  |  |  |  |  |
| TMB | 50 | 0.456 | 0.507 | 195 | 0.298 | 0.342 | 0.267 |
| Neoepitopes | 50 | 0.470 | 0.507 | 195 | 0.302 | 0.334 | 0.378 |
| TVB | 27 | 0.367 | 0.490 | 61 | 0.307 | 0.424 | 0.079 |
| NSCLC |  |  |  |  |  |  |  |
| TMB | 33 | 0.291 | 0.578 |  |  |  | 0.034 |
| Neoepitopes | 33 | 0.345 | 0.583 |  |  |  | 0.053 |
| RCC |  |  |  |  |  |  |  |
| TMB | 50 | 0.656 | 0.666 |  |  |  | 0.894 |
| Neoepitopes | 50 | 0.662 | 0.66 |  |  |  | 0.973 |
| TVB | 17 | 0.641 | 0.538 |  |  |  | 0.600 |

Coverage-adjusted SNV burden predicted response to immunotherapy better than overall TMB or any indel burden for all cancer types except RCC, for which coverage-adjusted burden of in-frame indels was the most predictive burden (see Supplementary Figure 8). Neoepitope burden alone predicted response to immunotherapy comparatively well as TMB, calculated using both raw and coverage-adjusted counts (see Figure 4). There was no difference in predictive capacity between Class I vs Class II epitope burdens (see Supplementary Figure 9). Similarly, incorporation of proteasomal cleavage, TAP transport, and cell surface presentation did not improve predictive capacity compared to TMB and neoepitope burden (see Supplementary Figure 10). We also weighted neoepitope burden by several criteria hypothesized to be related to increased immunogenicity, including: number of amino acid mismatches per peptide, number of MHC alleles predicted to bind each peptide, and number of TCGA-expressed transcripts of origin for the peptide (see Methods). In all cases, these weighted burdens yielded similar predictive capabilities to TMB or unadjusted neoepitope burden, though mismatch- and mismatch-by-allele-weighted neoepitope burdens incrementally improved predictive capacity for RCC patients, and allele-weighted neoepitope burden incrementally improved predictive capacity for NSCLC patients (see Figure 4). Interestingly, global assessment of HLA presentation (unique HLA allele count per patient) added slight predictive capacity to TMB in melanoma, RCC, and NSCLC patients (see Figure 4). However, the capacity for any of these metrics to predict patient-level immunotherapy response varied substantially by cancer type, with the highest predictive power for the NSCLC cohort, but a very limited predictive capability in melanoma, RCC, or when pooled across all cancer types (see Figure 4). Indeed, TMB as calculated by consensus variant calls predicts immunotherapy response more poorly than the experimental noise of the breadth of genomic coverage (Mbp) obtained via DNA sequencing in melanoma, RCC, and when pooled across cancer types (see Supplementary Figure 11).

**Figure 4:**
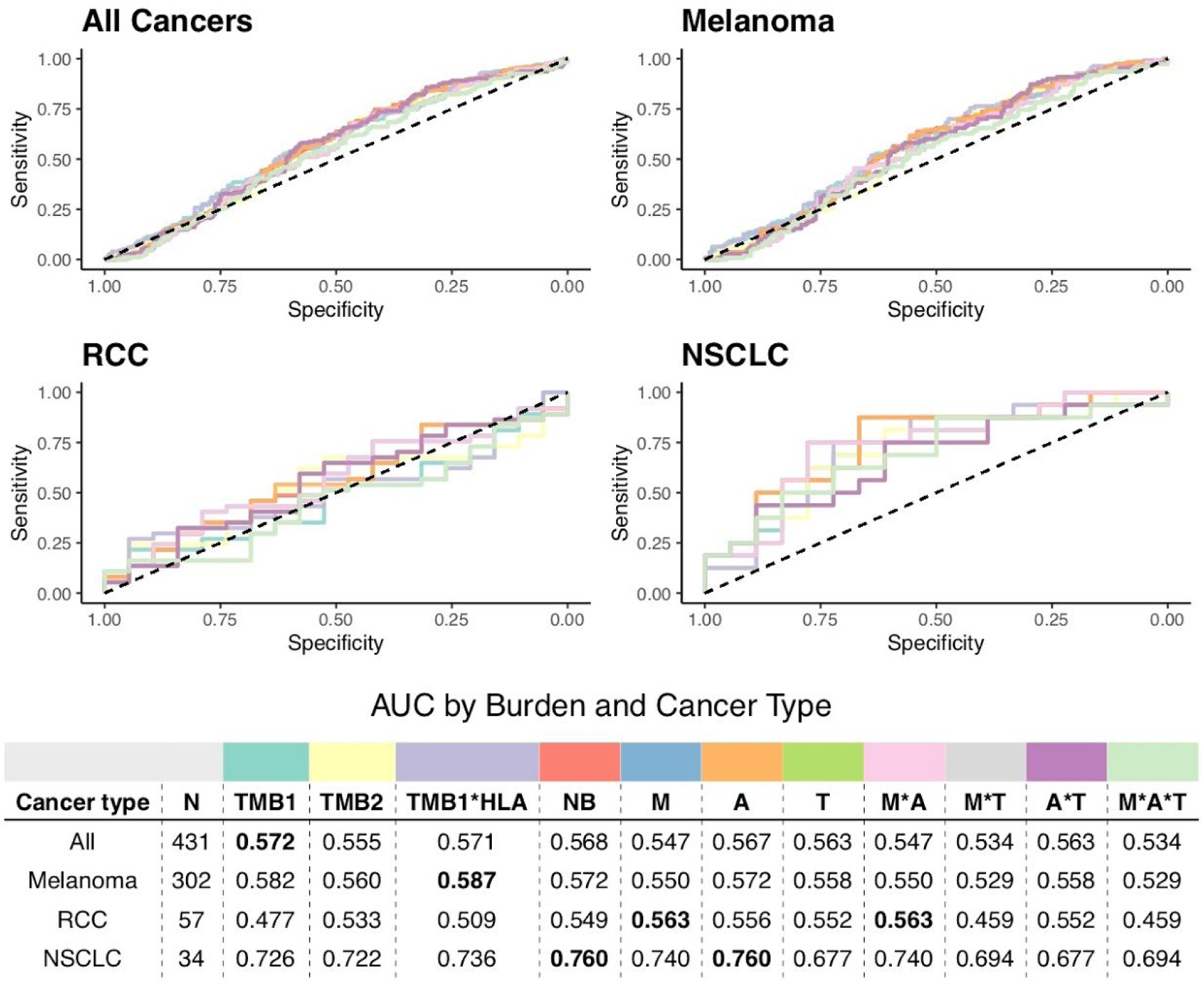
Receiver operating characteristic curves of predictive capacity of 11 different mutation/neoepitope burden metrics. The upper panels depict the true positive rate (sensitivity, y-axis) and false positive rate (1-specificity, x-axis) for each metric across all probability thresholds. The four panels represent models for four different cohorts based on different subsets of patients: All Cancers, which includes all patients, and Melanoma, RCC, and NSCLC, which include only melanoma, RCC, and NSCLC patients, respectively. The table in the lower panel reports the area-under-the-curve (AUC) for each metric (columns) applied to a different cancer cohort (rows), with colors above the methods indicating the color of the corresponding curve in the upper panels. TMB is used as a predictor in both raw (TMB1) and coverage-adjusted (TMB2) forms, as well as in a multiplicative combination with patient HLA allele count (TMB1*HLA). Neoepitope burden (NB) is used as a predictor in both raw and extended formats (see Methods). Extended neoepitope burden metrics include number of amino acid mismatches (M), number of HLA alleles predicted to bind each epitope (A), and number of transcripts expressing each epitope in TCGA (T), along with their multiplicative combinations. Bold-faced values indicate the best value for each cancer cohort.

For patients with tumor RNA-seq data, we also investigated how TVB and RNA-derived neoepitopes predicted response to immunotherapy (see Figure 5). We specifically considered tumor-specific junction burden, retained intron burden, retained intron neoepitope burden, and patient-specific expression-weighted neoepitope burdens (see Methods). As before, the vast majority of metrics (e.g. TMB, TVB) were all comparable in terms of predictive performance. However, considering these RNA-derived features did not increase predictive capacity over TMB, with the exception of the burden of tumor-specific splicing junctions, which yielded an increase in predictive performance for RCC patients (see Figure 5).

**Figure 5:**
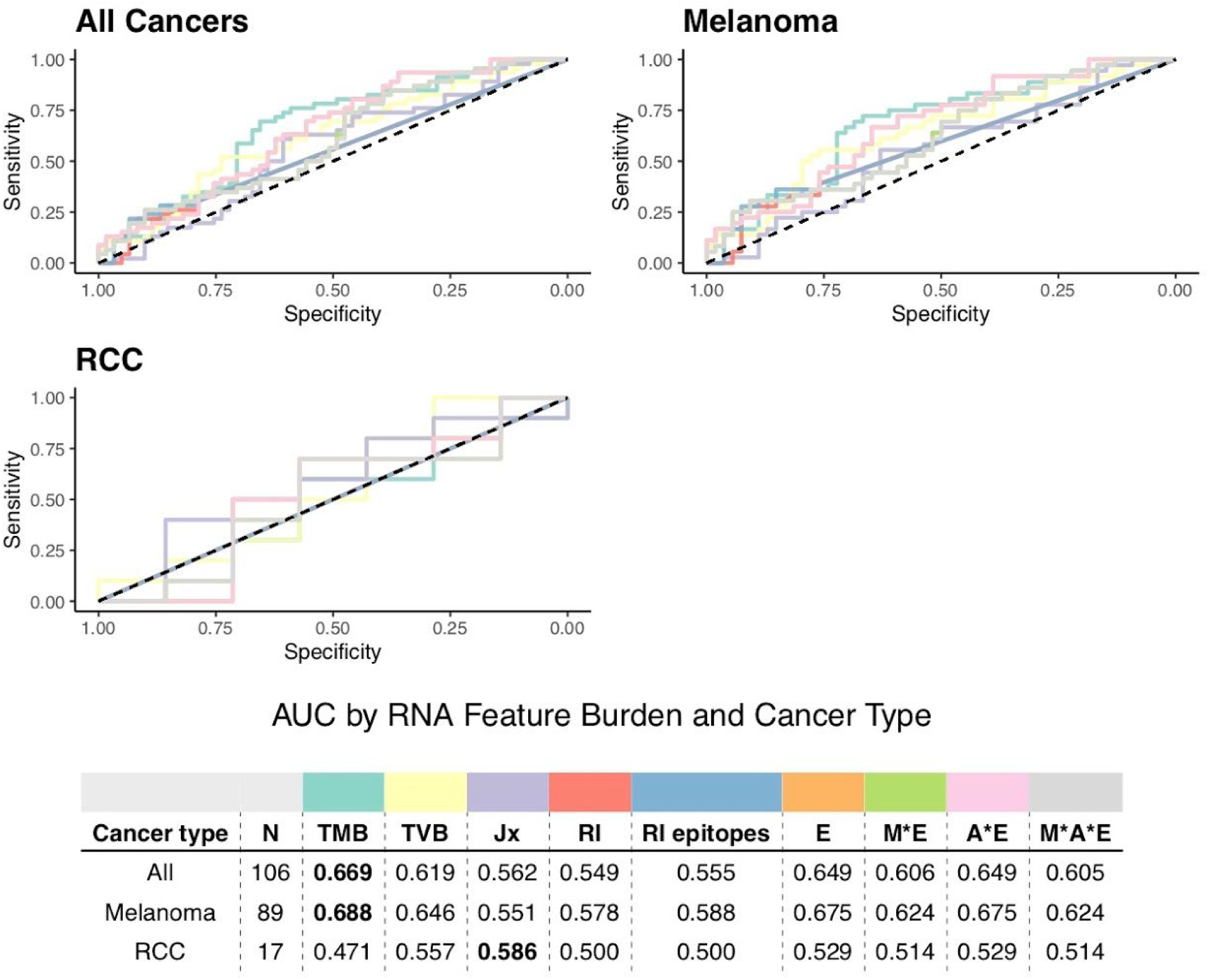
Receiver operating characteristic curves of predictive capacity of 9 different variant/neoepitope burden metrics. The upper panels depict the true positive rate (sensitivity, y-axis) and false positive rate (1-specificity, x-axis) for each metric across all probability thresholds. The three panels represent models for three different cohorts based on different subsets of patients: All Cancers, which includes all patients, and Melanoma, and RCC, which include only melanoma and RCC patients, respectively. The table in the lower panel reports the area-under-the-curve (AUC) for each metric (columns) applied to a different cancer cohort (rows), with colors above the methods indicating the color of the corresponding curve in the upper panels. TMB and TVB are used as predictors in the raw formats. Jx represents the number of tumor-specific junctions per patient, and RI represents the number of retained introns per patient, with RI epitopes representing neoepitopes derived from those retained introns. Neoepitope burden is used as predictor in its RNA-feature-extended formats (see Methods). Extended neoepitope burden metrics include number of expressed transcripts for each epitope (E), number of amino acid mismatches (M), number of HLA alleles predicted to bind each epitope (A), and number of transcripts expressing each epitope in TCGA (T), along with their multiplicative combinations.

Using an established threshold for identifying tumors with “high” TMB, namely TMB that exceeds the disease-matched 80th percentile (2), we investigated the metric’s capacity to predict overall survival in the context of immune checkpoint blockade therapy. While not statistically significant (p > 0.05, based on Cox proportional hazard modeling), we saw a clear trend towards improved overall survival among individuals with renal cell carcinoma and a high TMB (Figure 6A). Additionally, model comparisons using different TMB percentile cutoffs suggest that differences in overall survival for high and low TMB groups may be threshold dependent and alter model significance (see Supplementary Figure 12). Notably, the lack of a significant TMB effect may be due to insufficient sample size as the number of patients qualifying as high TMB decreases steadily with increasing threshold. In contrast, the same trend is not seen between TMB and overall survival among a separate cohort of patients (TCGA) in the absence of immunotherapy (Figure 6A). We also observed no differences in survival among individuals with metastatic melanoma (Figure 6B). In both cases, TVB and neoepitope burden demonstrate comparable capacities to stratify overall survival as TMB (Supplementary Figures 13 and 14).

**Figure 6:**
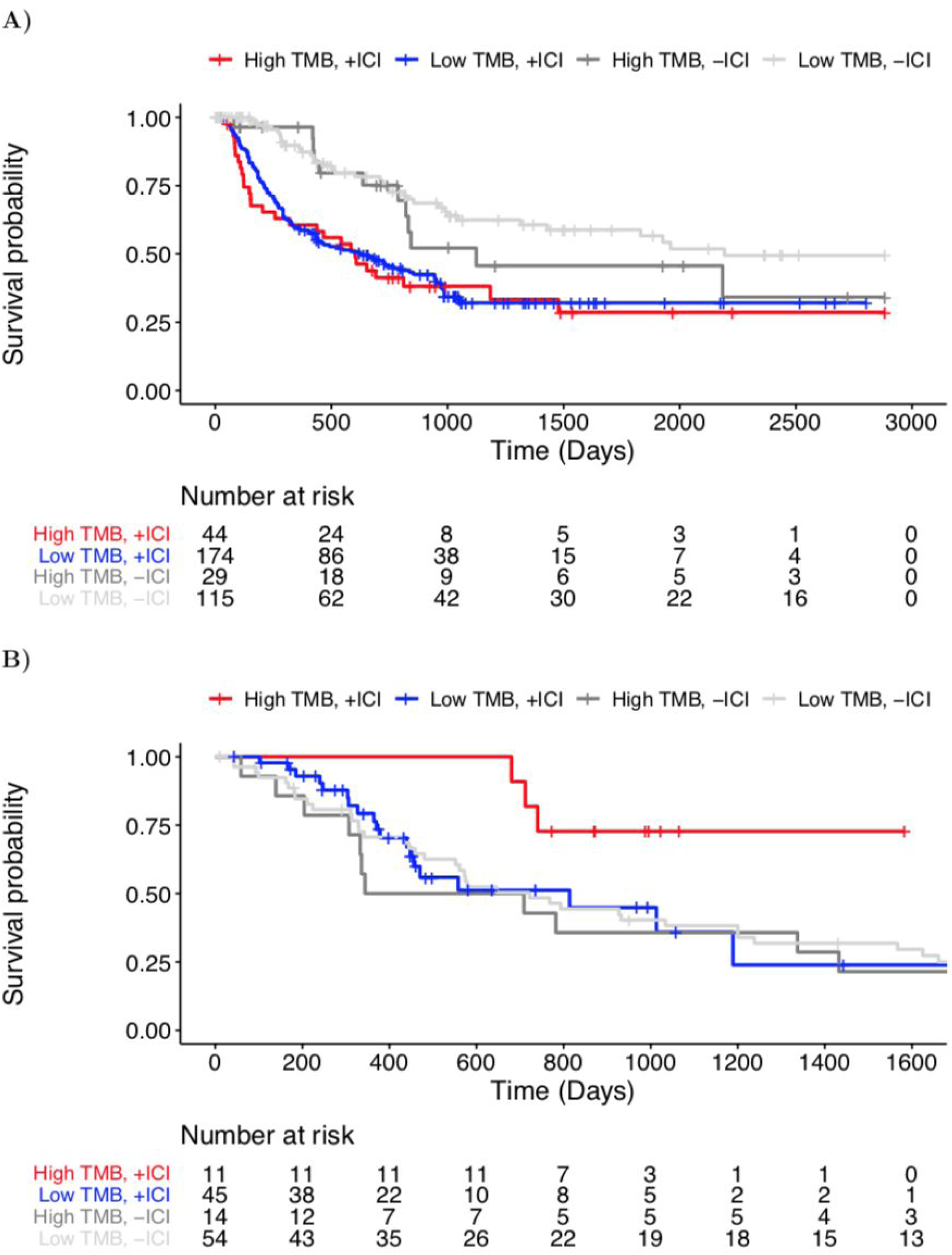
Overall survival among cancer patients with high and low TMB. A) Kaplan-meier curves for immunotherapy-treated (+ICI) and immunotherapy-naive (-ICI) Stage III-IV melanoma patients with high TMB (>80th percentile) are shown in red, and dark gray, respectively, while immunotherapy-treated (+ICI) and immunotherapy-naive (-ICI) patients with low TMB (≤80^th^ percentile) are shown in blue and light gray, respectively. The underlying table corresponds to the number of patients at risk of death at each timepoint. Note: TCGA SKCM patient data (-ICI) is censored at 2,885 days (maximal follow-up in immunotherapy-treated cohort) to emphasize comparable time-scales. B) Kaplan-meier curves for the immunotherapy-treated (+ICI) and immunotherapy-naive (-ICI) metastatic (Stage IV) renal cell carcinoma patients with high TMB (>80th percentile) are shown in red, and dark gray, respectively, while immunotherapy-treated (+ICI) and immunotherapy-naive (-ICI) patients with low TMB (≤80th percentile) are shown in blue and light gray, respectively. The underlying table corresponds to the number of patients at risk of death at each timepoint. Note: TCGA KIRC patient data is censored at 1,724 days (maximal follow-up in immunotherapy-treated cohort) to emphasize comparable time-scales.

### Metric instability of tumor variant and neoepitope burdens

We find, however, that TMB is not robust across variant calling methods. TMB as reported by individual variant calling tools was moderately similar to that reported by consensus calls (see Supplementary Figure 15), but variability in per-caller TMB increased with increasing number of variants (see Supplementary Figure 16). Additionally, the difference in TMB between the highest and lowest counts from individual callers per patient (median difference of 1,840 variants per patient) reflects a substantial fraction of the overall TMB, accounting for a median 59.3% of the value in the metric overall (see Methods).

We also compared tumor mutational burden as reported by the authors of the original manuscripts from which our cohort originated with our standardized consensus approach. While author-reported and consensus values for TMB were significantly correlated (Pearson’s product-moment correlation of 0.35, p = 1.99×10^−7^; see Supplementary Figure 17), we note that author-reported values have a universally higher predictive capacity than we observe using consensus data (Supplementary Figure 18). We also find important discrepancies in per-patient classification. Approximately 26.2% of patients are incongruously determined to be TMB “high” or “low” (using a TMB threshold >80th percentile as per (2)), however as many as 42.23% of patients may be dubiously classified using alternative thresholds (e.g. 29th-65th percentiles; see Supplementary Figure 19). Consensus and author-reported nonsynonymous mutation burdens exhibited a similar extent of correlation as well as per-patient instability of classification (Pearson’s product-moment correlation of 0.58, p < 2.2×10^−16^; see Supplementary Figures 18 and 20). The correlation between consensus-derived neoepitope burden and that reported by the original manuscripts was weak and not statistically significant (Pearson’s product-moment correlation of 0.026, p = 0.70; see Supplementary Figure 21). Moreover, we find that response hazard ratios are not stable based on TMB thresholds, a phenomenon especially dramatic in the RCC cohort (see Supplementary Figure 12), and consistent with prior findings that a single TMB threshold is inappropriate to apply across different cancer types (2).

Finally, we find that the predictive performance of TMB is sensitive to the method(s) used to perform variant calling (see Figure 7). Note that the same phenomenon holds true for raw TMB counts (see Supplementary Figure 22). While outside the scope of the current manuscript, note also that the identity of resulting neoepitopes is also highly sensitive to variant calling method (see Supplementary Figure 23).

**Figure 7:**
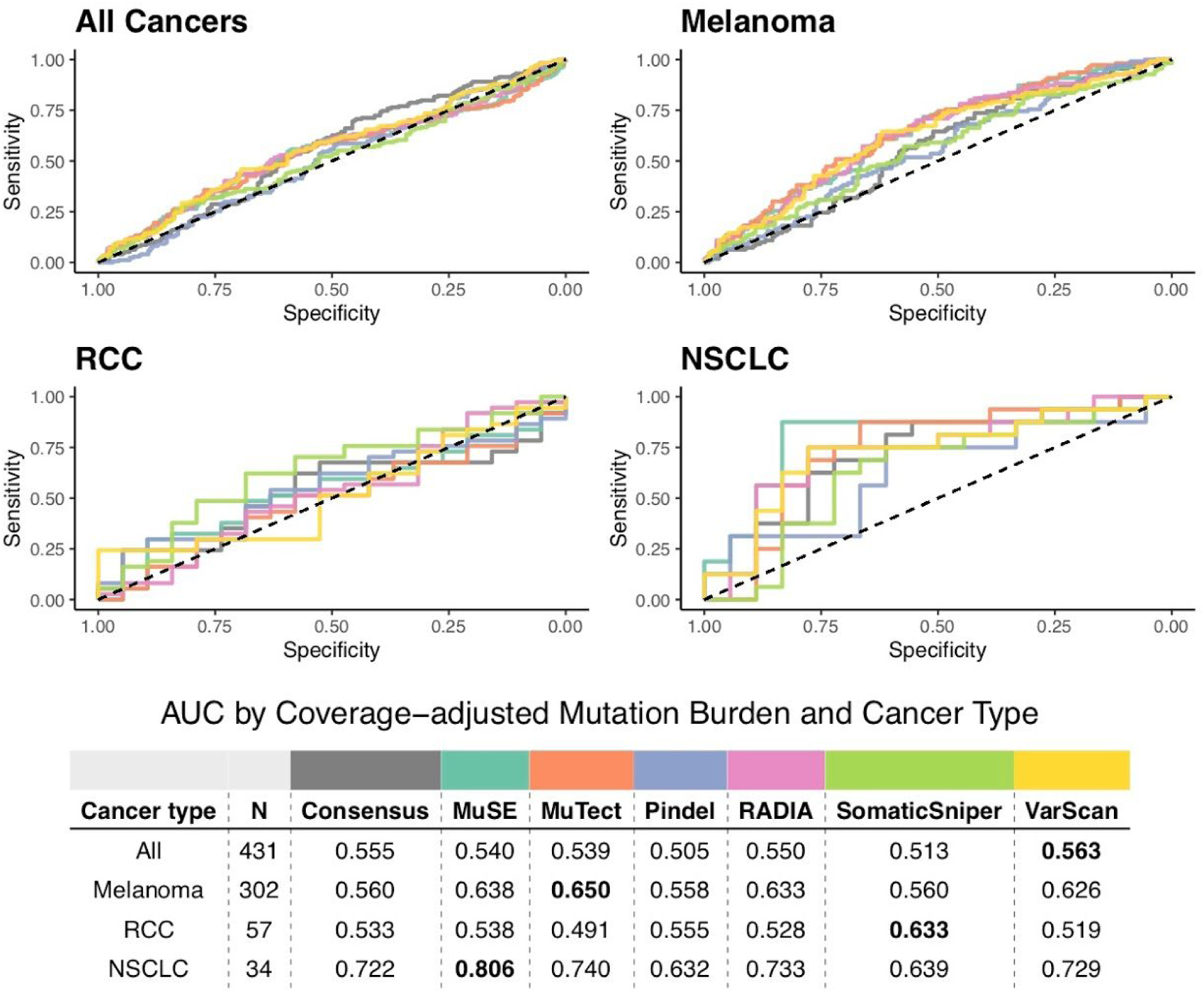
Receiver operating characteristic curves of predictive capacity of coverage-adjusted TMB from 7 different variant calling methods. The upper panels depict the true positive rate (sensitivity, y-axis) and false positive rate (1-specificity, x-axis) for each method across all probability thresholds. The four panels represent models for four different cohorts based on different subsets of patients: All Cancers, which includes all patients, and Melanoma, RCC, and NSCLC, which include only melanoma, RCC, and NSCLC, respectively. The table in the lower panel reports the area-under-the-curve (AUC) for each method (columns) applied to a different cancer cohort (rows), with colors above the methods indicating the color of the corresponding curve in the upper panels. TMB as determined by consensus calling (see Methods) is compared to the individual variant calling tools used in consensus calling. RCC=renal cell carcinoma, NSCLC=non-small cell lung cancer.

## DISCUSSION

To the best of our knowledge, this is the first study to evaluate TMB and correlated downstream metrics such as neoepitope burden from whole exome sequencing data using a gold standard ensemble approach (31,61) applied to a meta-cohort of immunotherapy-treated cancer patients across multiple studies and disease types. This study also introduces the concept of tumor variant burden, incorporating potential RNA-derived sources of variants where available, and is the first study to estimate immunotherapy response rate as a function of TMB, TVB, and neoepitope burden. Moreover, this study is the first to quantitatively evaluate the stability of TMB as a metric, and the first to directly compare the predictive capacities of multiple TMB and related metrics.

Ultimately, we show that TMB is a dubious predictor of immunotherapy response, with substantial caveats regarding: 1) predictive capacity differences among different cancer types, with RCC being no better than random chance, 2) sensitivity of TMB and downstream metrics to variant calling methodology, and 3) stability of TMB thresholds and their ability to classify patients in a population. This suggests that the prospective clinical utilization of TMB is likely subject to many of these same issues, and may result in unintended harms, whether due to omission of therapy for individuals with “low” TMB who might nonetheless benefit, or due to increased risk of toxicity in a “high” TMB population subject to overuse of immunotherapy. Indeed, a recent study of metastatic melanoma patients (62) found significantly different burdens of nonsynonymous mutations between disease subgroups, but not between progressors/responders, highlighting the instability of this metric.

With rare exception, we find no added predictive benefit to evaluating more complex bulk metrics downstream of TMB. Akin to prior observations, incorporation of HLA genotype diversity adds slightly to the predictive capacity of TMB (63). Given the added technical effort and costs required to perform these analyses, we conclude that TMB is likely the optimal bulk assessment of tumor variation among those tested, though inclusion of HLA diversity data may marginally improve estimates. However, such bulk measurements neglect the potential importance of individual cancer neoantigens, which recent evidence suggests may be the driving force behind response to cancer immunotherapy by eliciting tumor-antigen-specific T cell responses (64).

This study has several limitations. First, numerous sampling based assays have also been used to assess TMB (e.g. (2,65,66)), however, we did not evaluate these data in this study, instead focusing on whole exome sequencing data as the prevailing gold standard for accurate mutational assessment. Note that these targeted assays would not enable incorporation of HLA allelic diversity data into a predictive model. Note also that there is wide variability among TMB assay design, analysis, and performance, with the potential for overestimation of TMB when using gene-targeted assays (67). Ultimately, along with the substantial variability among widely-used targeted assays (11), and the futility of expecting universal adoption of a single technique, this study highlights the need for increased standardization of TMB interpretation, a subject of active pursuit by the TMB Harmonization Project (68). Second, we did not compare TMB in this dataset with other potential predictors of immunotherapy response (e.g. based on gene expression (69) or copy number instability (70)), however it is possible that TMB could be synergistic with such orthogonal metrics. Third, by virtue of the retrospective nature of these data and limited availability of whole exome sequencing cohorts, this study cannot be assumed to translate to emerging immunotherapies and instead is interpretable exclusively for αPD1 and αCTLA4 therapy.

While this study is consistent with multiple prior reports demonstrating the importance of TMB in predicting immunotherapy response (e.g. (2,71)), the caveats raised herein are of high concern for the field overall. Our collective emphasis on TMB is understandable given its relative ease of quantification using various techniques, however it is indeed a dubious and indirect predictor. Tumors with higher TMB have been hypothesized to have more neoantigens that can be recognized by the immune system in response to checkpoint inhibition, yet the data presented here and data previously published (2) support the use of substantially different “absolute” TMB thresholds for immunotherapy response prediction across different diseases. Further, evidence suggests that other genomic factors, such as tumor purity and clonal heterogeneity, may further modulate the relationship between TMB and immunotherapy response (62,72). This suggests an added layer of as-of-yet undefined complexity not captured in the current bulk metrics, and likely related to disease-specific biology.

## CONCLUSIONS

In conclusion, we find sufficient cause to suggest that the predictive clinical value of TMB should not be overstated or oversimplified. While it is readily quantified, TMB is at best a limited surrogate biomarker of immunotherapy response. The data confirms TMB as a reasonable predictor in non-small cell lung cancer, and a weak predictor in melanoma. The data does not support TMB in isolation as a predictive biomarker for RCC, though it may be feasibly combined with HLA allelic diversity to achieve marginal performance.

## LIST OF ABBREVIATIONS

αCTLA4: Anti-cytotoxic T-lymphocyte-associated protein 4
αPD1: Anti-programmed cell death protein 1
AUC: Area Under the Curve
COAD: Colon Adenocarcinoma
GATK: Genome Analysis Toolkit
GTEx: Genotype Tissue Expression
HLA: Human Leukocyte Antigen
Indel: Insertion or deletion
KIRC: Kidney Renal Clear Cell Carcinoma
kma: Keep Me Around
LUAD: Lung Adenocarcinoma
LUSC: Lung Squamous Cell Carcinoma
MHC: Major Histocompatiblity Complex
MMR: Mismatch Repair
NSCLC: Non Small Cell Lung Cancer
PRAD: Prostate Adenocarcinoma
RCC: Renal Cell Carcinoma
RI: Retained intron
RNA-seq: RNA sequencing
ROC: Receiver Operating Characteristic
SKCM: Skin Cutaneous Melanoma
SRA: Sequence Read Archive
TCGA: The Cancer Genome Atlas
THCA: Thyroid carcinoma
TMB: Tumor Mutational Burden
TPM: Transcripts Per Million
TVB: Tumor Variant Burden
UCEC: Uterine Corpus Endometrial Carcinoma
VCF: Variant Call Format
WES: Whole Exome Sequencing

## DECLARATIONS

**Ethics approval and consent to participate**: Not applicable

**Consent for publication**: Not applicable

**Availability of data and materials:** Data used in these analyses is available on the Sequence Read Archive under accessions PRJNA278450, PRJNA293912, PRJNA305077, PRJNA306070, PRJNA307199, PRJNA312948, PRJNA324705, PRJNA343789, PRJNA357321, PRJNA369259, PRJNA414014, PRJNA420786, and PRJNA82745 and on the European Genome-Phenome Archive under the accession EGAD00001004352. Data from Le et al. (9) and Graff et al. (17) are available from those authors upon reasonable request. The results of our analyses of these data are available as Supplemental Tables 2-7 in Additional File 1. We have created a GitHub repository with instructions for reproducing results, available at https://github.com/pdxgx/immunorx_response_pipeline.

## Disclaimer

The contents do not represent the views of the U.S. Department of Veterans Affairs or the United States Government.

## Author’s contributions

Study and conceptual design was by MAW, AN, and RFT. Data analysis was by MAW, BRW, and JKD. Manuscript writing was predominantly by MAW, AN, and RFT, with contributions from BRW and JKD. Final manuscript review and approval was by MAW, BRW, JKD, AN, and RFT.

## Acknowledgments

Thank you to Adam Struck and Kyle Ellrott for their support implementing somatic variant calling pipelines. Thank you to Julie Graff and Rachel Slottke for providing prostate cancer cohort data.

## SUPPLEMENTARY DATA

**Supplementary Table 1:** Summary of patients samples used for analysis. Publicly available WES data from 12 studies was used to determine TMB (see Materials and Methods). We summarize the study which produced each data set, the cancer types represented, and the number of patients/tumor samples sequenced. Studies that had complementary RNA sequencing reads available for at least a subset of patients are indicated by an asterisk in the “Reference” column, and the number of samples with complementary RNA sequencing data are indicated in parentheses in the “Number of patients” and “Number of tumor samples” columns if different than the number of samples with WES.

| Cancer type | Number of patients | Number of tumor samples | Reference |
| --- | --- | --- | --- |
| Melanoma | 15 | 15 | Amaria et al. |
| Melanoma | 3 | 6 | Carreno et al.* |
| Melanoma | 17 | 17 | Eroglu et al. |
| Melanoma | 16 | 16 | Gao et al. |
| Prostate | 10 | 10 | Graff et al. |
| Melanoma | 38 (27) | 39 (28) | Hugo et al.* |
| Colon, endometrial, thyroid | 28 | 29 | Le et al. |
| RCC | 57 (17) | 58 (17) | Miao et al. |
| NSCLC | 34 | 34 | Rizvi et al. |
| Melanoma | 35 | 53 | Roh et al. |
| Melanoma | 64 (20) | 64 (20) | Snyder et al.* |
| Melanoma | 110 (40) | 110 (40) | Van Allen et al.* |
| Melanoma | 4 | 6 | Zaretsky et al. |

Supplementary Tables 2-7 summarize the results of our analyses of genomic data, and they are available in Additional File 1. Column descriptions for these tables follow. For privacy, we have only included summary data in these tables and not identities of predicted variants or their resulting neopeptides.

Supplementary Table 2: Summary of patient clinical and DNA variant data. The “Patient” column indicates the patient identifier, the “Tumor_ID” column indicates the tumor sample identifier(s), and the “Normal_ID” column indicates the normal sample identifier(s); for patients with more than one tumor sample, median values are presented in subsequent columns of the table where relevant. The “Disease” column indicates the cancer type of the patient, and the “Study” column indicates the first author of the study from which the genomic data originated. The “Coverage” column indicates the Mbp of genome covered by at least 6 sequencing reads. The “Total_mutations” column indicates the total number of somatic DNA variants, the “SNVs” column indicates the number of single nucleotide variants, the “Inframe_insertions” column indicates the number of in-frame insertion variants, the “Inframe_deletions” column indicates the number of in-frame deletion variants, the “Frameshift_insertions” column indicates the number of frameshifting insertion variants, the “Frameshift_deletions” column indicates the number of frameshifting deletion variants, and the “Nonsynonymous_SNVs” indicates the number of single nucleotide variants resulting in a protein-level change; all of these variants burdens are the result of consensus variant calling from variants predicted by MuSE, MuTect, Pindel, RADIA, SomaticSniper, and VarScan 2. For each individual variant caller, there are four columns summarizing their total variant counts, SNVs, insertions, and deletions; for example, the “Muse_variants” column indicates the total number of variants predicted by MuSE, the “Muse_SNVs” column indicates the number of single nucleotide variants predicted by MuSE, the “Muse_deletions” column indicates the number of deletions predicted by MuSE, and the “Muse_insertions” column indicates the number of insertions predicted by MuSE. The “Tumor_HLA1_count” column indicates the number of unique MHC Class I alleles for the patient, and the “Tumor_HLA2_count” column indicates the number of unique MHC Class II alleles for the patient. The “MSI_status” column indicates whether the patient was MSI-high as predicted by mSINGs (binary). The “Cancer_stage” column indicates the cancer stage of the patient at the time of the original study. The “aPD1_treatment” column indicates whether the patient was given anti-PD1 treatment (binary) over the course of the original study, the “aPDL1_treatment” column indicates whether the patient was given anti-PDL1 treatment (binary) over the course of the original study, the “aCTLA4_treatment” column indicates whether the patient was given anti-CTLA4 treatment (binary) over the course of the original study, and the “Other_treatment” column indicates whether the patient was given any other treatment (binary) over the course of the original study. The “aCTLA4_response” column indicates whether the patient responded to anti-CTLA4 treatment (binary) over the course of the original study, the “aPD1_response” column indicates whether the patient responded to anti-PD1 treatment (binary) over the course of the original study, and the “Combined_response” column indicates whether the patient responded to any treatment (binary) over the course of the original study. The “PFS” column indicates the number of days the patient went without progression of disease over the course of the original study, the “OS” column indicates the number of days the patient was alive over the course of the original study, and the “Censoring_days” column indicates the number of days the patient was monitored over the course of the original study. The “Vital_status” column indicates whether the patient was deceased (binary) at the end of the original study, the “OS_event” column indicates whether the patient had a disease-related death (binary) during the course of the original study for patients, the “PFS_event” column indicates whether the patient experienced disease progression (binary) during the course of the original study, and the “Censoring_status” column indicates whether the progression/survival data for the patient was censored (binary) at the end of the original study. The “Original_nonsynonymous_mutations” column indicates the burden of nonsynonymous somatic mutations reported by the authors of the original study, the “Original_total_mutations” column indicates the burden of total somatic mutations reported by the authors of the original study, and the “Original_neoantigens” column represents the burden of neoantigens reported by the authors of the original study. The “Total_unphased_neoepitopes” column indicates the number of neopeptide sequences predicted from consensus somatic variants if haplotype phasing of variants is not considered, and the “Total_comprehensive_neoepitopes” column indicates the number of neopeptide sequences predicted from consensus somatic variants if haplotype phasing of variants is considered. The “MHCnuggets_eps” column indicates the number of consensus-calling-derived neoepitopes predicted to bind to one or more of a patient’s MHC Class I or Class II alleles as predicted by MHCnuggets, the “MHCnuggets_ClassI_eps” column indicates the number of consensus-calling-derived neoepitopes predicted to bind to one or more of a patient’s MHC Class I alleles as predicted by MHCnuggets, and the “MHCnuggets_ClassII_eps” column indicates the number of consensus-calling-derived neoepitopes predicted to bind to one or more of a patient’s MHC Class II alleles as predicted by MHCnuggets. The “Manuscript_binding_eps” column indicates the number of consensus-calling-derived neoepitopes predicted using the same peptide size and HLA allele restrictions as the authors of the original study (see Methods).

Supplementary Table 3: Per-patient summary of driver variants and neoepitopes. The “Patient” column indicates the patient identifier, and the “Tumor_ID” column indicates the tumor sample identifier(s); for patients with more than one tumor sample, median values are presented in subsequent columns of the table. The “Total_clinvar_variants” column indicates the number of variants annotated as “pathogenic” or “likely pathogenic” in cancer by ClinVar, and the “Total_clinvar_neopeptides” column indicates the number of neopeptide sequences derived from these variants. The “Presented_clinvar_variants” column indicates the number of variants in the “Total_clinvar_variants” column that have at least one neopeptide predicted to bind to one or more of the patient’s MHC Class I or Class II alleles. The “Presented_clinvar_epitopes” column indicates the number of neopeptides in the “Total_clinvar_neopeptides” column that are predicted to bind to one or more of the patient’s MHC Class I or Class II alleles.

Supplementary Table 4: Modified neoepitope burdens. The “Patient” column indicates the patient identifier, and the “Tumor_ID” column indicates the tumor sample identifier(s); for patients with more than one tumor sample, median values are presented in subsequent columns of the table. The “Epitope_by_mismatch_burden” column sums for each neopeptide the number of amino acids changes it contains. The “Epitope_by_allele_burden” column sums for each neopeptide the number of MHC Class I or Class II alleles predicted to bind to that neopeptide. The “Epitope_by_expression_burden” column sums for each neopeptide the number of transcripts expressed in patient RNA-seq data (where available) that would give rise to that neopeptide. The “Epitope_by_TCGA_burden” sums for each neopeptide the number of transcripts expressed in the matched TCGA cancer type (see Methods) that would give rise to that neopeptide. The “Epitope_by_mismatch_and_allele_burden” column sums for each neopeptide the number of amino acid changes it contains multiplied by the number of MHC Class I or Class II alleles predicted to bind to that neopeptide. The “Epitope_by_mismatch_and_expression_burden” column sums for each neopeptide the number of amino acid changes it contains multiplied by the number of patient-expressed transcripts that would give rise to that neopeptide. The “Epitope_by_mismatch_and_TCGA_burden” column sums for each neopeptide the number of amino acid changes it contains multiplied by the number of transcripts expressed in the matched TCGA cancer type that would give rise to that neopeptide. The “Epitope_by_allele_and_expression_burden” column sums for each neopeptide the number of MHC Class I or Class II alleles predicted to bind to that neopeptide multiplied by the number of patient-expressed transcripts that would give rise to that neopeptide. The “Epitope_by_allele_and_TCGA_burden” column sums for each neopeptide the number of MHC Class I or Class II alleles predicted to bind to that neopeptide multiplied by the number of transcripts expressed in the matched TCGA cancer type that would give rise to that neopeptide. The “Epitope_by_mismatches_alleles_and_expression_burden” column sums for each neopeptide the number of amino acid changes it contains multiplied by the number of MHC Class I or Class II alleles predicted to bind to that neopeptide multiplied by the number of patient-expressed transcripts that would give rise to that neopeptide. The “Epitope_by_mismatches_alleles_and_TCGA_burden” column sums for each neopeptide the number of amino acid changes it contains multiplied by the number of MHC Class I or Class II alleles predicted to bind to that neopeptide multiplied by the number of transcripts expressed in the matched TCGA cancer type that would give rise to that neopeptide.

Supplementary Table 5: Tumor-specific splice junction burdens. The “Patient” column indicates the patient identifier, and the “Tumor_ID” column indicates the tumor sample identifier(s). The “Jx_burden” column indicates the number of tumor-specific splice junctions per patient; for patients with more than one tumor sample, median values are presented.

Supplementary Table 6: Tumor-specific retained intron and retained intron epitope burdens. The “Patient” column indicates the patient identifier, and the “Tumor_ID” column indicates the tumor sample identifier(s); for patients with more than one tumor sample, median values are presented in subsequent columns of the table. The “Intron_burden” column indicates the number of tumor-specific retained introns per patient. The “Intron_epitope_burden” indicates the number of neopeptide sequences associated with tumor-specific retained introns for that patient, and the “Binding_intron_epitope_burden” indicates the number of these peptides predicted to bind to one or more of the patient’s MHC Class I or Class II alleles.

Supplementary Table 7: Processed neoepitope burdens. The “Patient” column indicates the patient identifier, and the “Tumor_ID” column indicates the tumor sample identifier(s). The “NetCTLpan_epitopes” column represents the number of neopeptides predicted by NetCTLpan to be proteasomally processed, TAP transported, and cell-surface presented for at least one patient MHC Class I allele; for patients with more than one tumor sample, the median value across all samples is presented in this column.

**Supplementary Figure 1:**
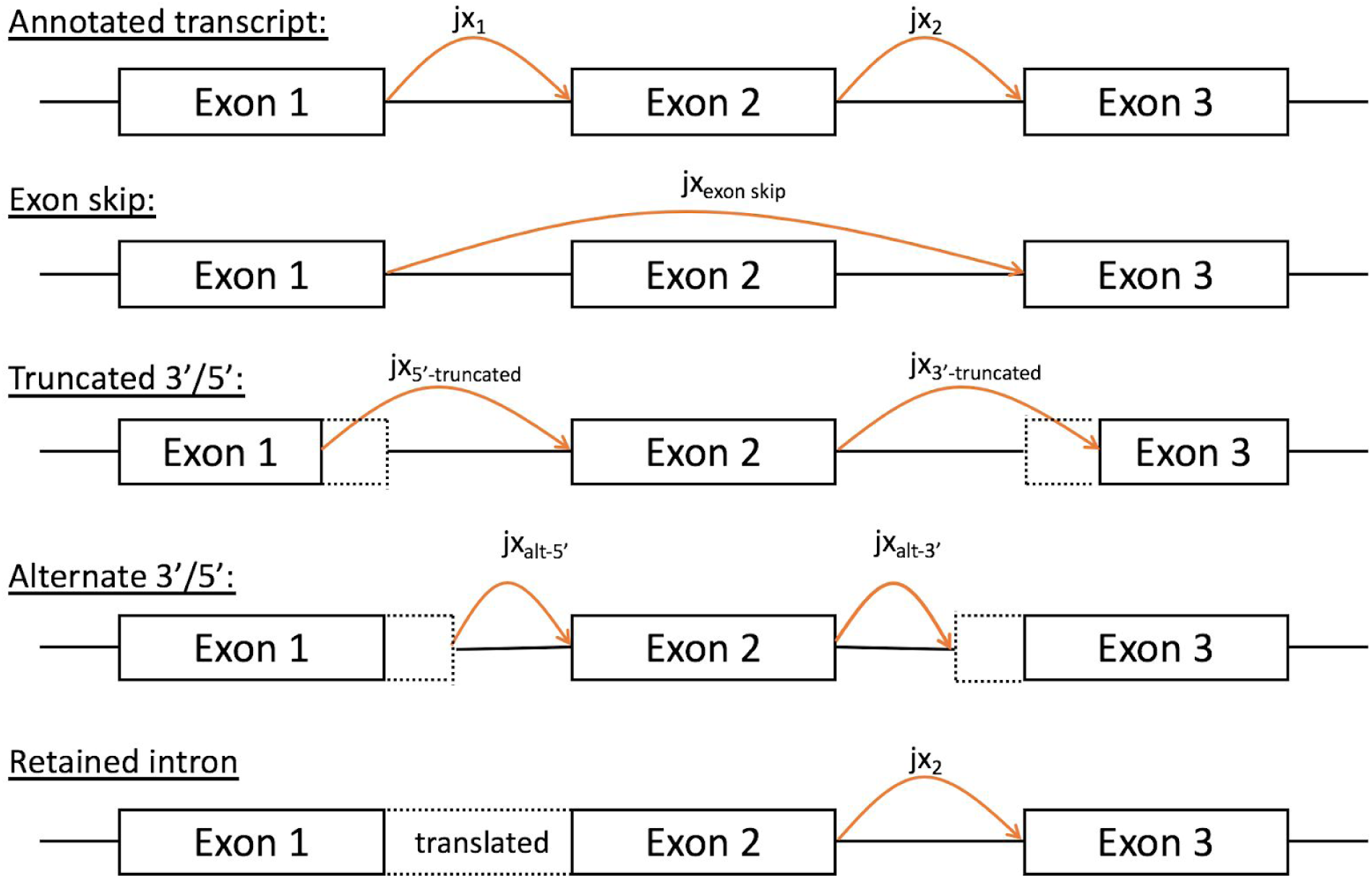
Visual depiction of potential splice variants captured. The top row shows the annotated “normal” splicing for a simulated gene with 3 exons; this splicing is represented by junctions (jxs) 1 and 2. A potential exon skip is represented on row 2, where exon 2 is skipped. Possible alternate 5’ and 3’ splice sites are shown in rows 3 and 4, and a retained intron between exons 1 and 2 in row 5.

**Supplementary Figure 2:**
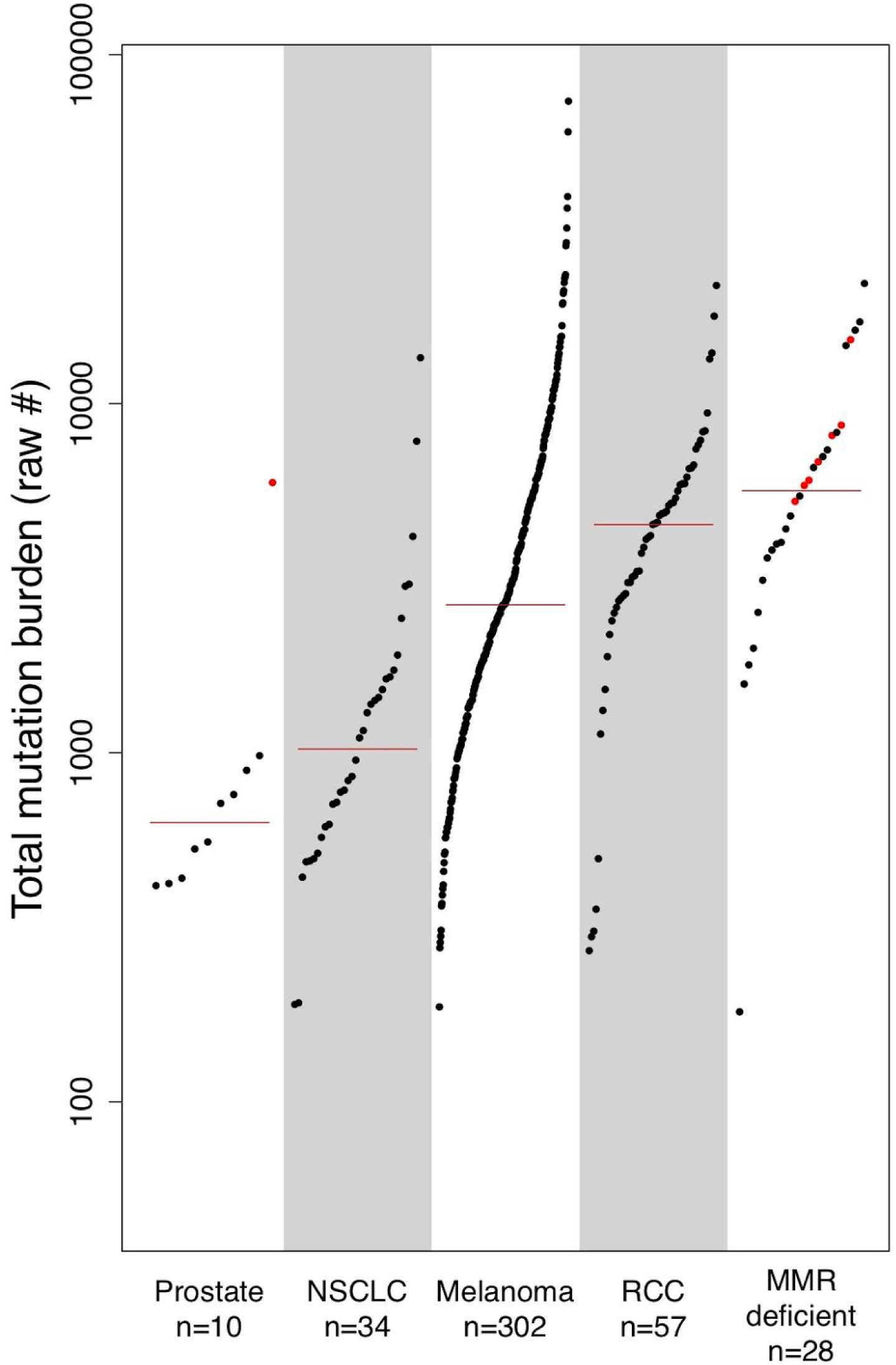
Per-patient distribution of raw mutation burdens across 7 cancer types. The raw number of somatic DNA variants per patient are shown along the y-axis, with each dot representing an individual cancer patient (cancer types shown along the x-axis). Note that MMR-deficient cancers here represent a cohort of 3 different cancer types including colon, endometrial, and thyroid with evidence of mismatch repair deficiency as determined by polymerase chain reaction or immunohistochemistry (9). Red colored dots correspond to patients with microsatellite instability as determined by mSINGS (see Methods). Abbreviations as follows: RCC=renal cell carcinoma, NSCLC=non-small cell lung cancer, MMR=mismatch repair.

**Supplementary Figure 3:**
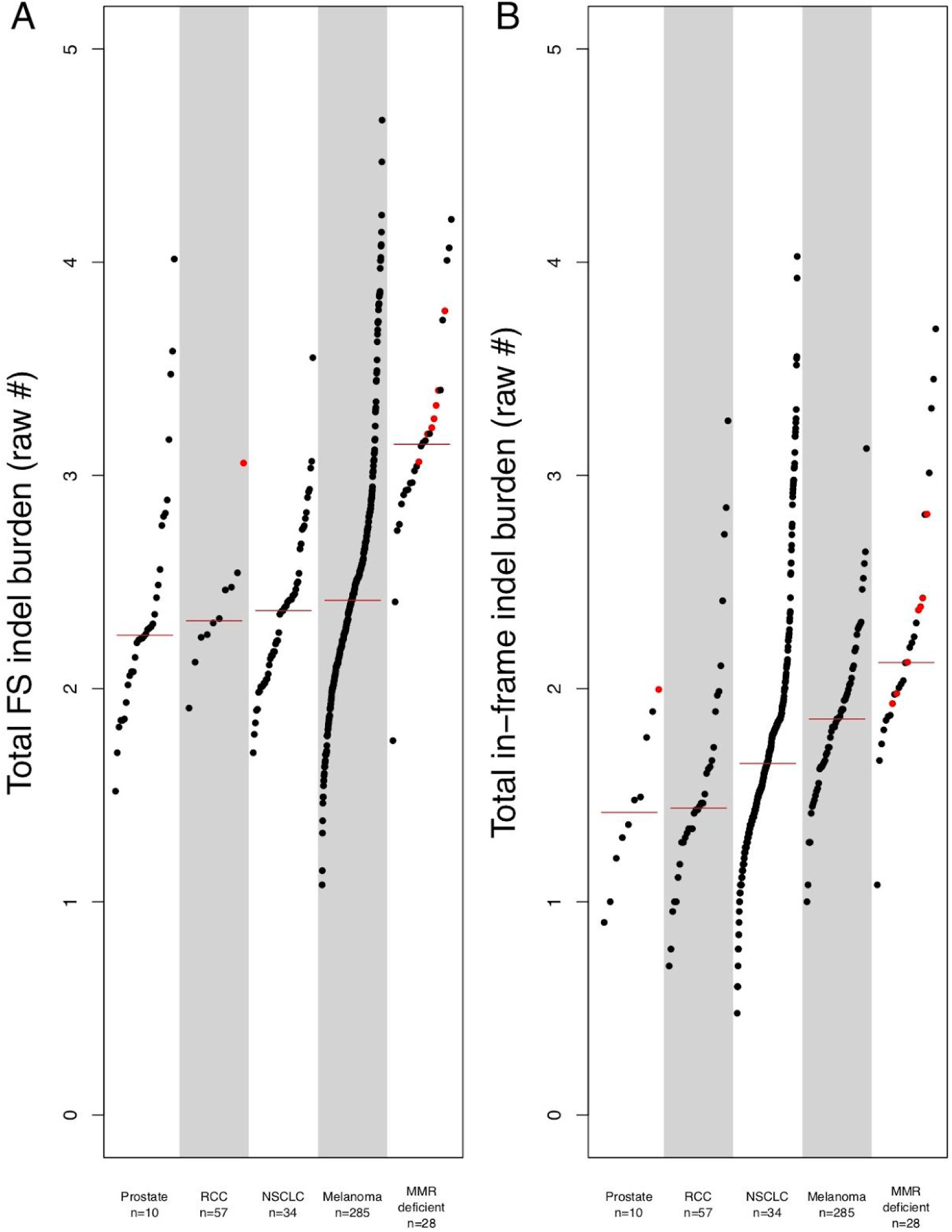
Per-patient distribution of insertion and deletion (indel) burdens across 7 cancer types. A) The number of somatic frameshift (FS) indels per patient are shown along the y-axis, with each dot representing an individual cancer patient (cancer types shown along the x-axis). Note that MMR-deficient cancers here represent a cohort of 3 different cancer types including colon, endometrial, and thyroid with evidence of mismatch repair deficiency as determined by polymerase chain reaction or immunohistochemistry (9). Red colored dots correspond to patients with microsatellite instability as determined by mSINGS (see Methods). B) The number of somatic in-frame indels per patient are shown along the y-axis, with each dot representing an individual cancer patient (cancer types shown along the x-axis). Abbreviations as follows: RCC=renal cell carcinoma, NSCLC=non-small cell lung cancer, MMR=mismatch repair.

**Supplementary Figure 4:**
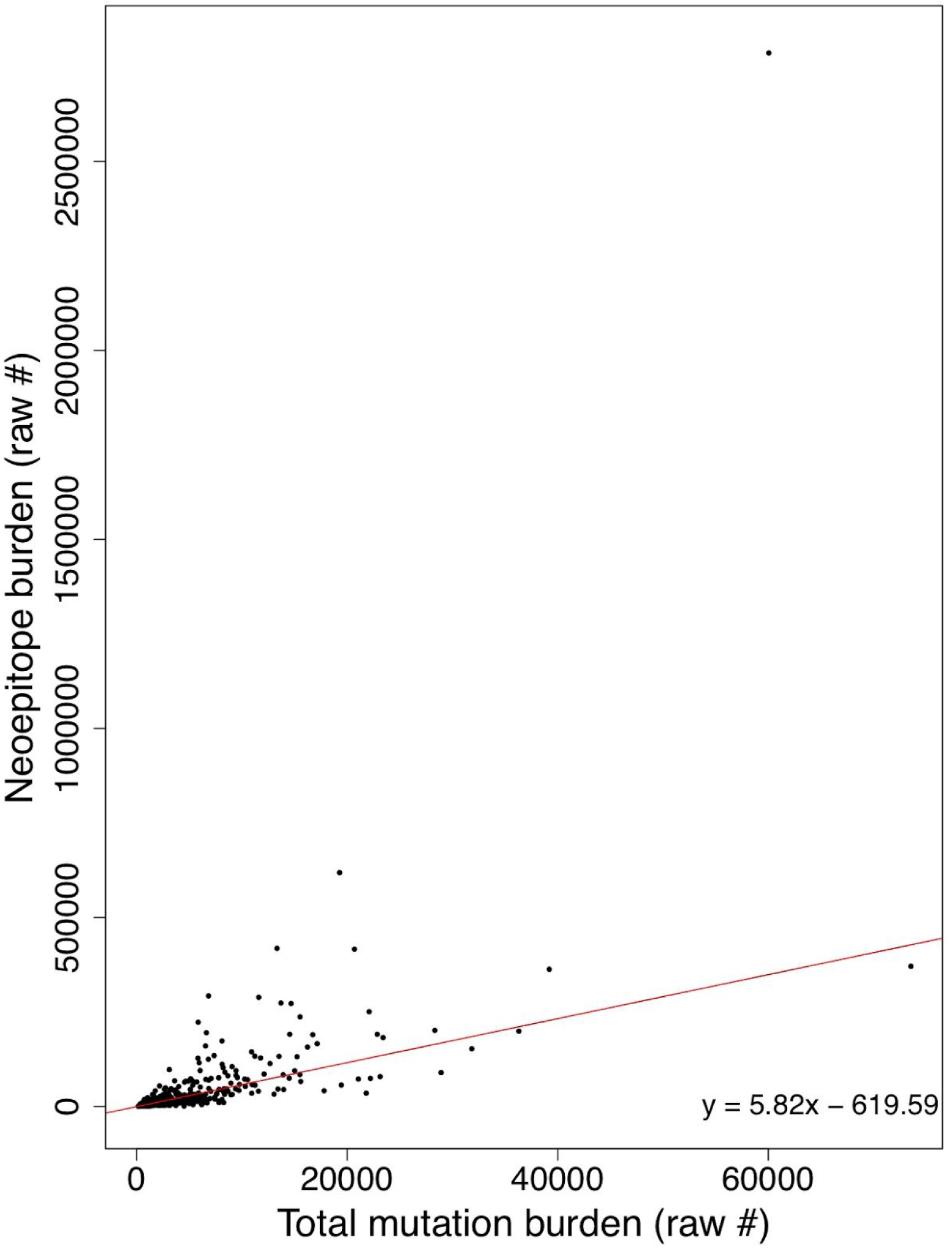
TMB correlates with neoepitope burden. Tumor mutational burden (x-axis) and neoepitope burden (y-axis) are strongly correlated (Pearson product-moment correlation of 0.63, p < 2.2×10^−16^). The best fit line as determined by linear regression is shown in red, with its equation in the bottom right corner.

**Supplementary Figure 5:**
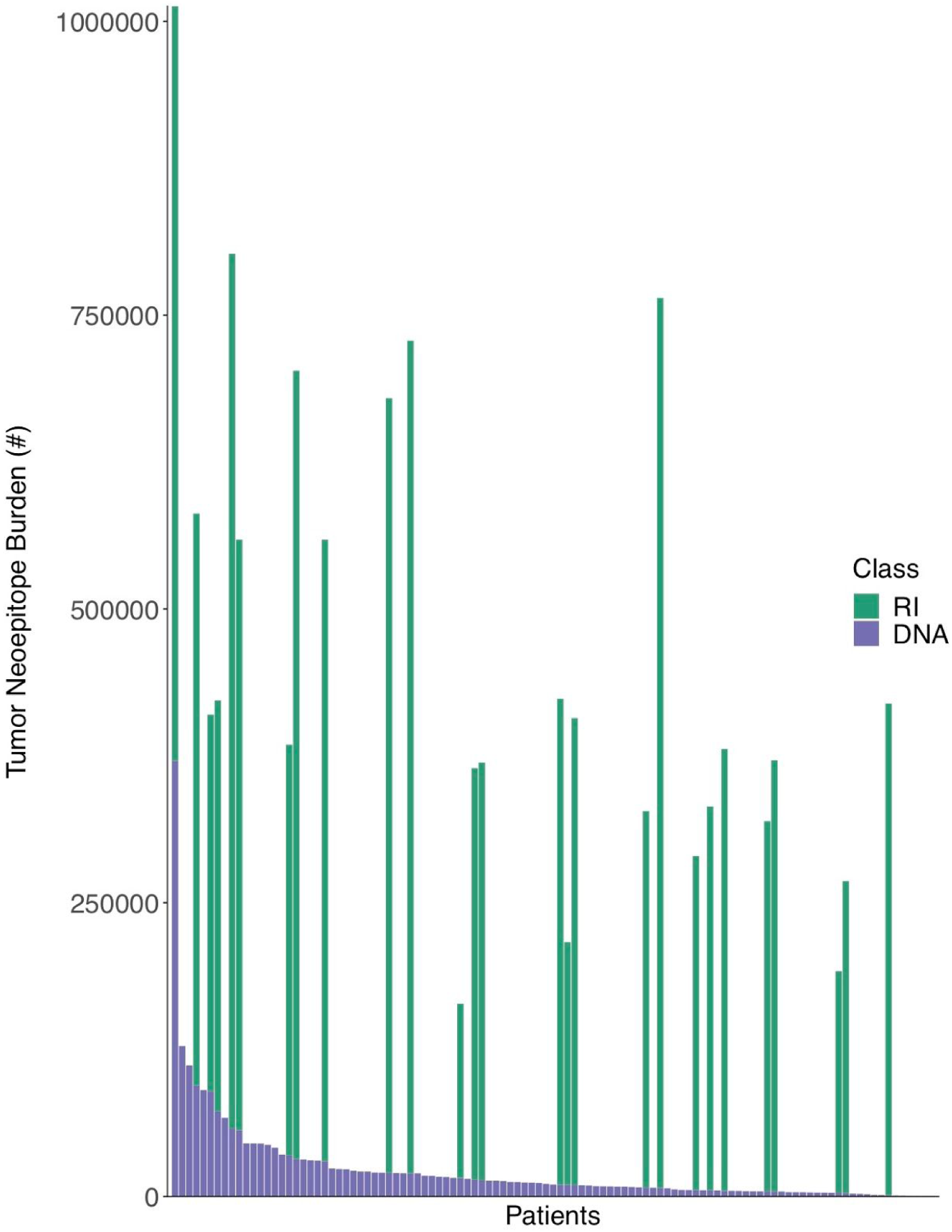
Per-patient distribution of overall tumor neoepitope burden and its components. The number of total tumor neoepitopes per patient is shown along the y-axis, with the numbers of neoepitopes derived from retained introns (RI) and somatic DNA variants (DNA) shown in green and purple, respectively. The data for each individual patient is displayed as stacked bars along the x-axis, sorted from left to right by the number of neoepitopes derived from somatic DNA variants (from highest to lowest).

**Supplementary Figure 6:**
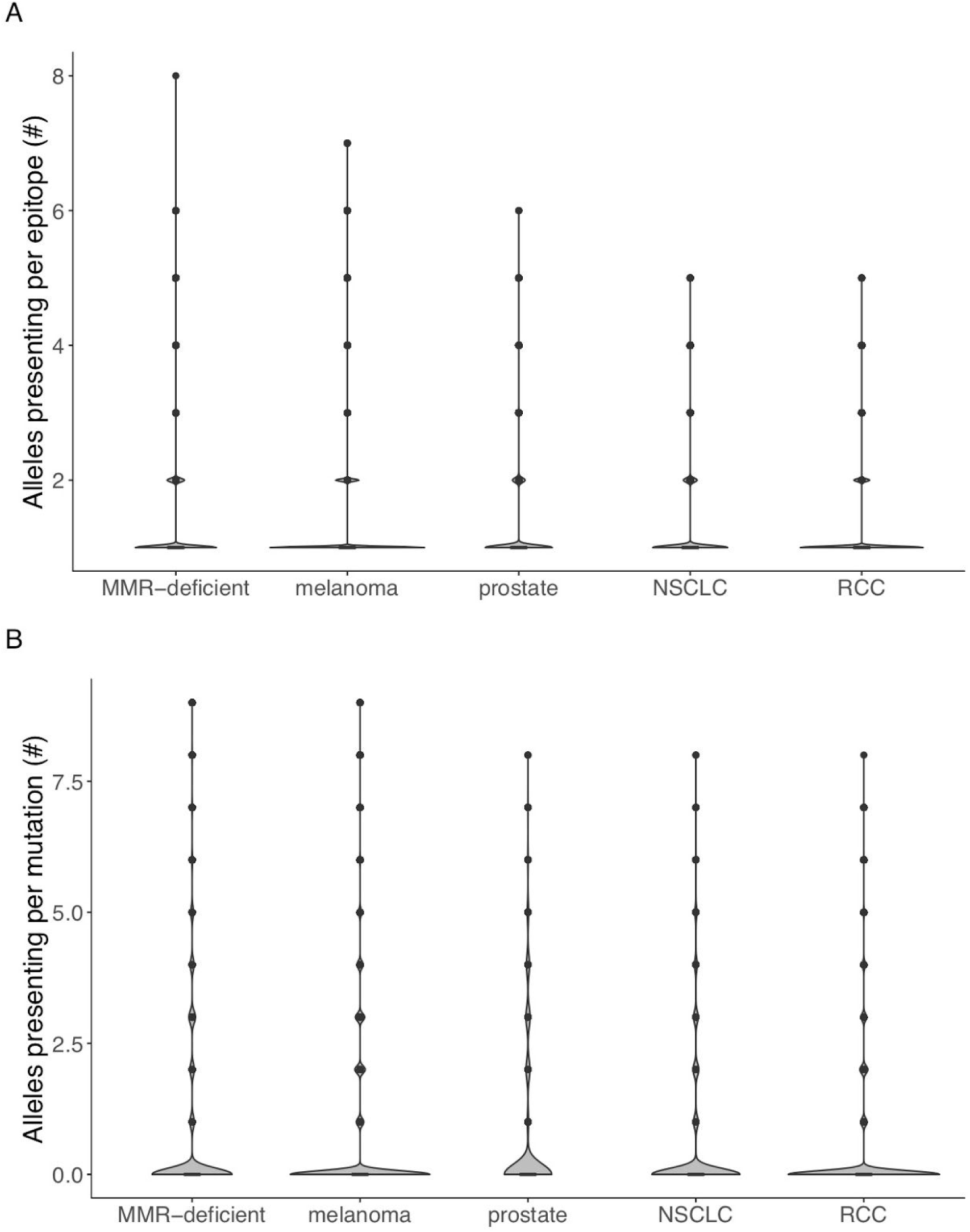
Robustness of putative neoepitope presentation among 5 different cancer groups. A) The number of unique patient-matched HLA alleles that are predicted to present an individual neoepitope is shown along the y-axis, with each violin plot distribution corresponding to a different cancer group along the x-axis, as labeled. Note that MMR-deficient cancers here represent a cohort of 3 different cancer types including colon, endometrial, and thyroid with evidence of mismatch repair deficiency as determined by polymerase chain reaction or immunohistochemistry (9). B) The total number of unique patient-matched HLA alleles that are predicted to present one or more neoepitopes arising from a single DNA mutation is shown along the y-axis, with each violin plot distribution corresponding to a different cancer group along the x-axis, as labeled. Note that the width of each violin plot at each point along the y-axis corresponds to the relative quantity of data points in that group for that value of the y-axis. Furthermore, the lower and upper borders of the box within each violin plot corresponds to the 25th and 75th percent quantiles of the dataset for that group, respectively, with the median value shown as a horizontal black line within the box. Note that a predicted HLA binding affinity threshold of ≤500nM was used in all cases (see Methods).

**Supplementary Figure 7:**
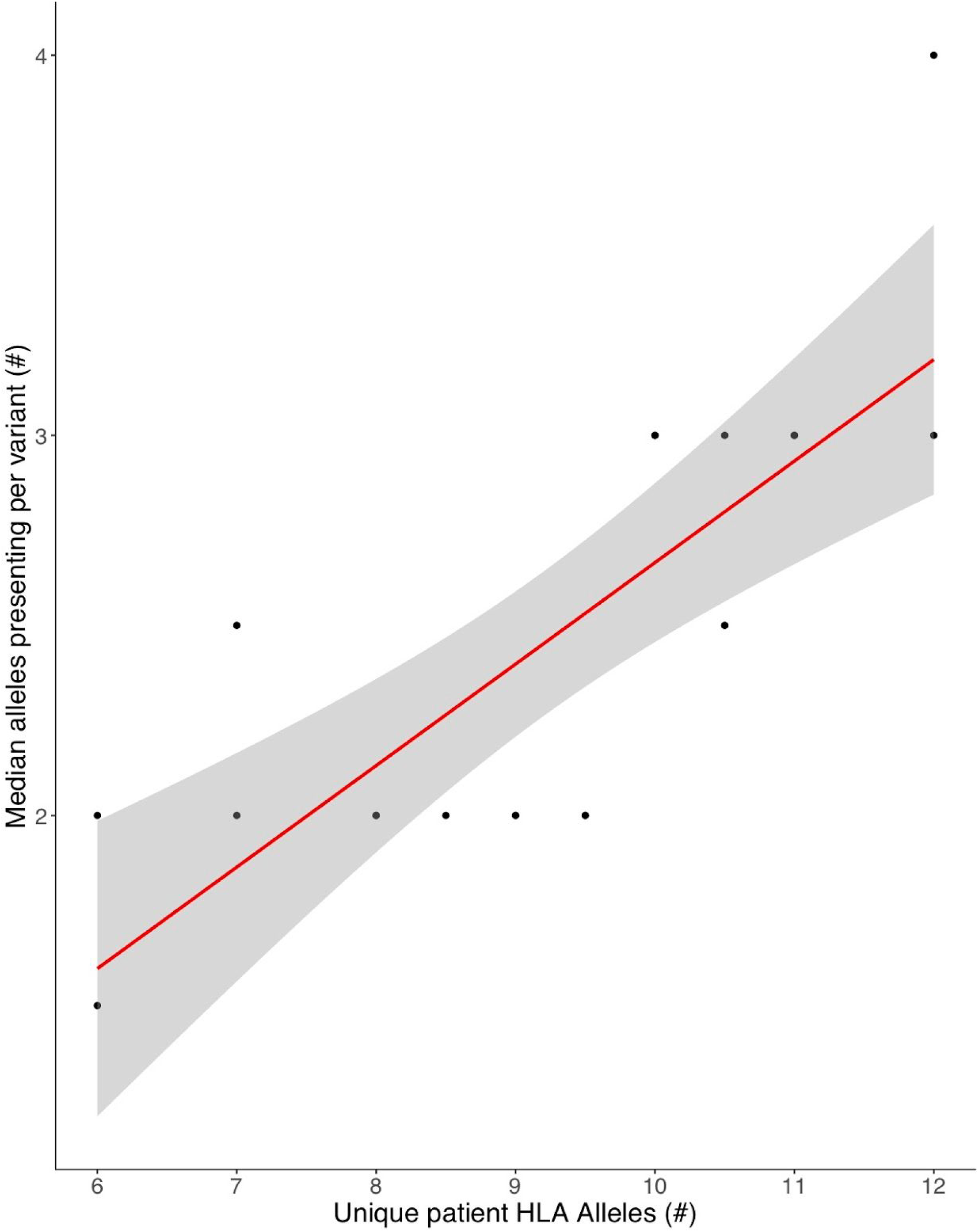
Robustness of putative neoepitope presentation. The median number of unique patient-matched HLA alleles that are predicted to present one or more neoepitopes arising from a single DNA mutation is shown along the y-axis, with the x-axis corresponding to patient-specific HLA heterozygosity (as the number of unique MHC I and II alleles per patient). Red curve denotes the best fit line based on linear regression, with surrounding gray shading denoting the 95% confidence interval. Note that a predicted HLA binding affinity threshold of ≤500nM was used in all cases (see Methods).

**Supplementary Figure 8:**
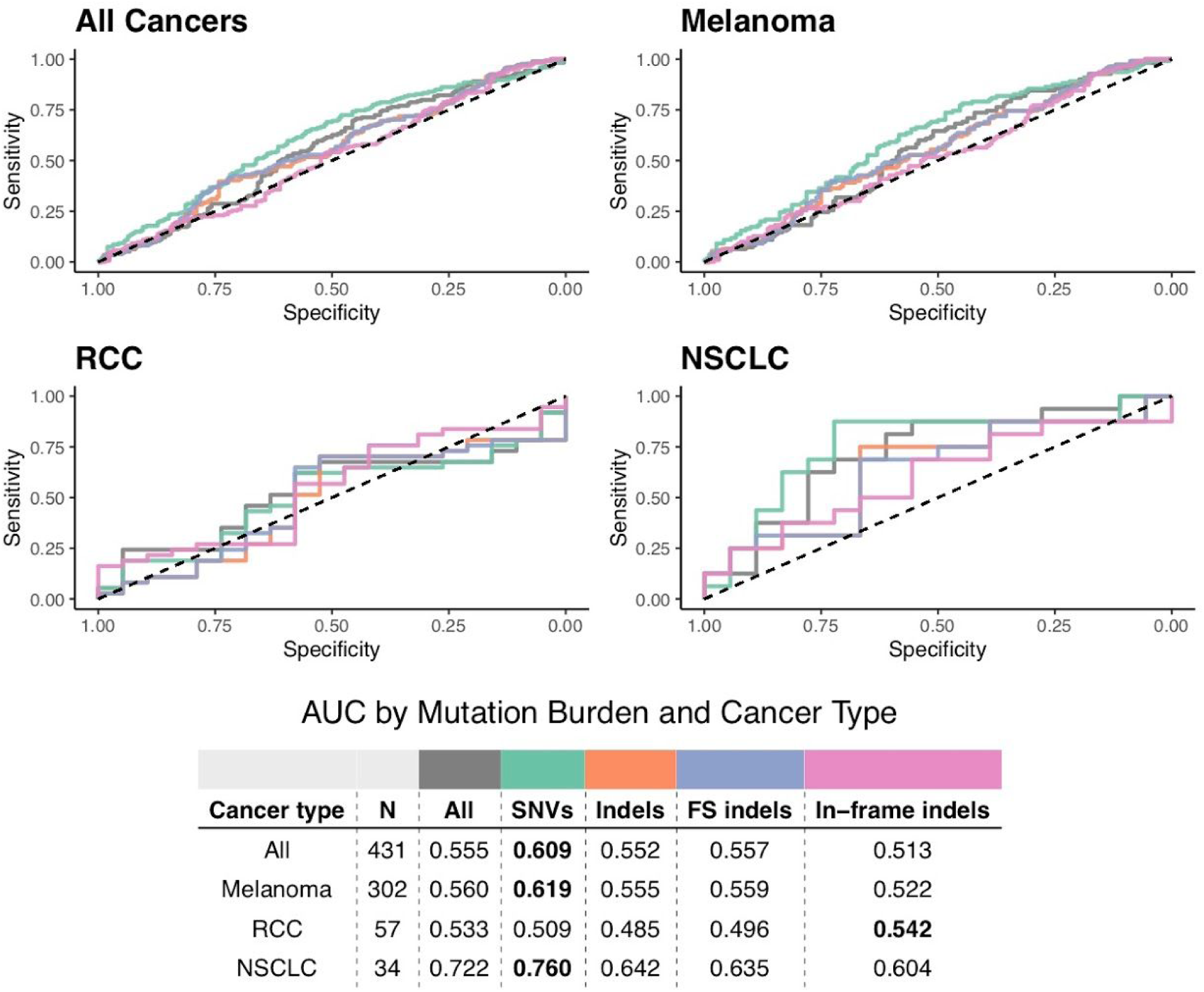
Receiver operating characteristic curves of predictive capacity of 5 different coverage-adjusted variant burden metrics. The upper panels depict the true positive rate (sensitivity, y-axis) and false positive rate (1-specificity, x-axis) for each metric across all probability thresholds. The three panels represent models for three different cohorts based on different subsets of patients: All Cancers, which includes all patients, and Melanoma, and RCC, which include only melanoma and RCC patients, respectively. The table in the lower panel reports the area-under-the-curve (AUC) for each metric (columns) applied to a different cancer cohort (rows), with colors above the methods indicating the color of the corresponding curve in the upper panels. All represents all DNA variants (SNVs and indels of all types), SNVs includes all single nucleotide variants, Indels includes all insertion/deletion variants, FS indels includes all frameshifting insertions and deletions, and In-frame indels includes all in-frame insertions and deletions.

**Supplementary Figure 9:**
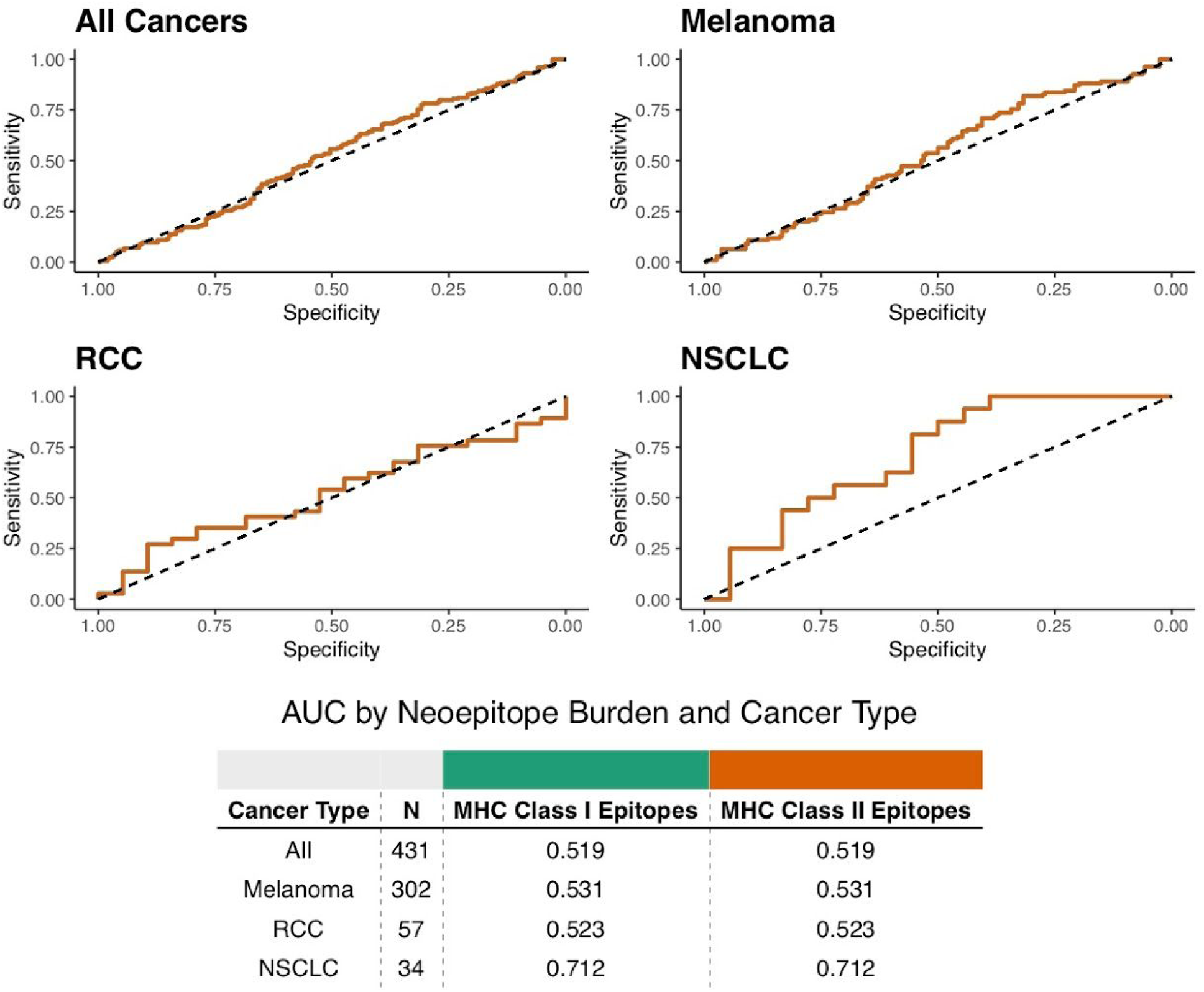
Receiver operating characteristic curves of predictive capacity of MHC Class I vs. MHC Class II neoepitope burdens. The upper panels depict the true positive rate (sensitivity, y-axis) and false positive rate (1-specificity, x-axis) for each metric across all probability thresholds. The three panels represent models for three different cohorts based on different subsets of patients: All Cancers, which includes all patients, and Melanoma, and RCC, which include only melanoma and RCC patients, respectively. The table in the lower panel reports the area-under-the-curve (AUC) for each metric (columns) applied to a different cancer cohort (rows), with colors above the methods indicating the color of the corresponding curve in the upper panels.

**Supplementary Figure 10:**
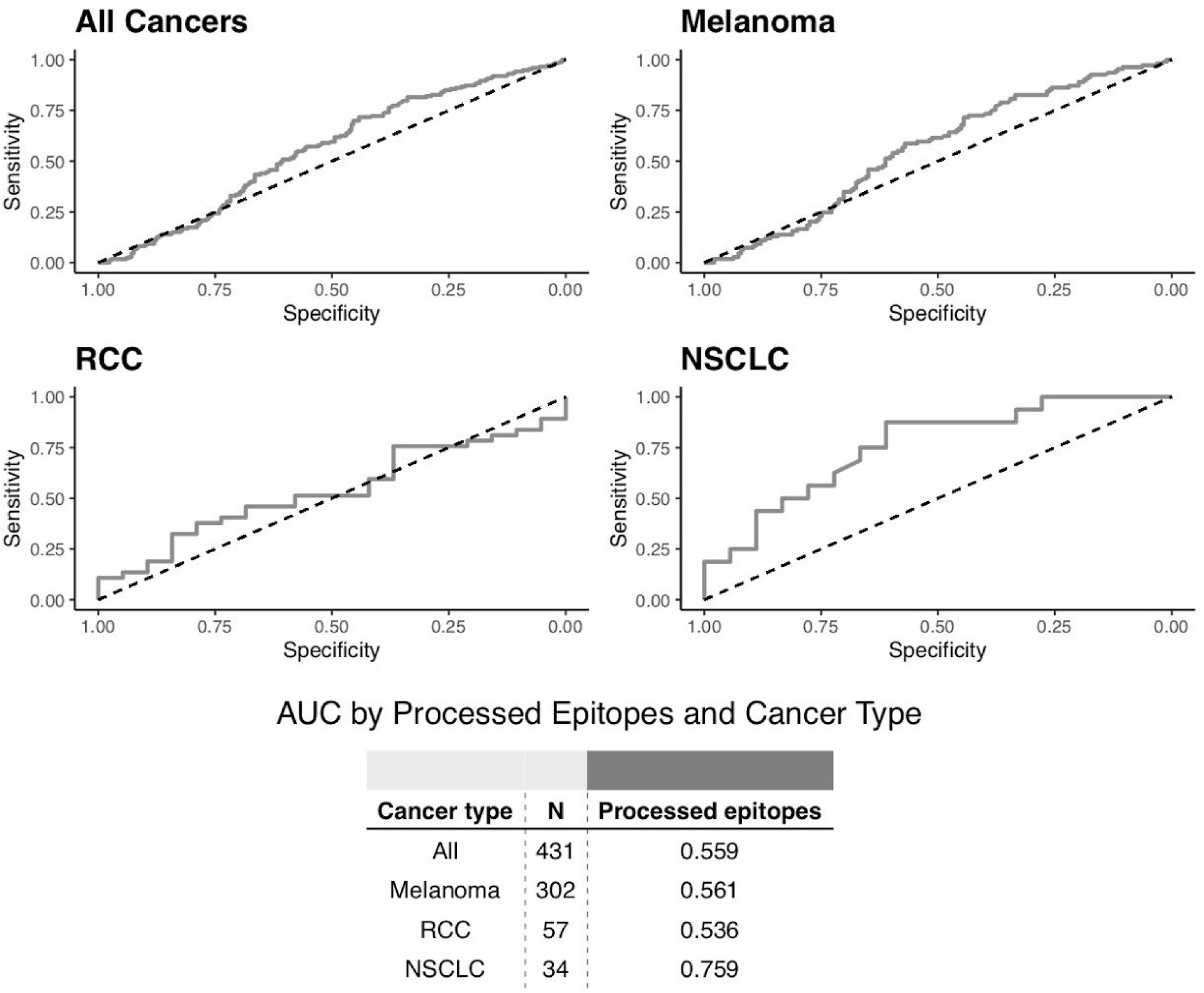
Receiver operating characteristic curves of predictive capacity of processed neoepitope burden. The upper panels depict the true positive rate (sensitivity, y-axis) and false positive rate (1-specificity, x-axis) for genomic coverage across all probability thresholds. The four panels represent models for four different cohorts based on different subsets of patients: All Cancers, which includes all patients, and Melanoma, RCC, and NSCLC, which include only melanoma, RCC, and NSCLC patients, respectively. The table in the lower panel reports the area-under-the-curve (AUC) for coverage (right column) applied to a different cancer cohort (rows). RCC=renal cell carcinoma, NSCLC=non-small cell lung cancer.

**Supplementary Figure 11:**
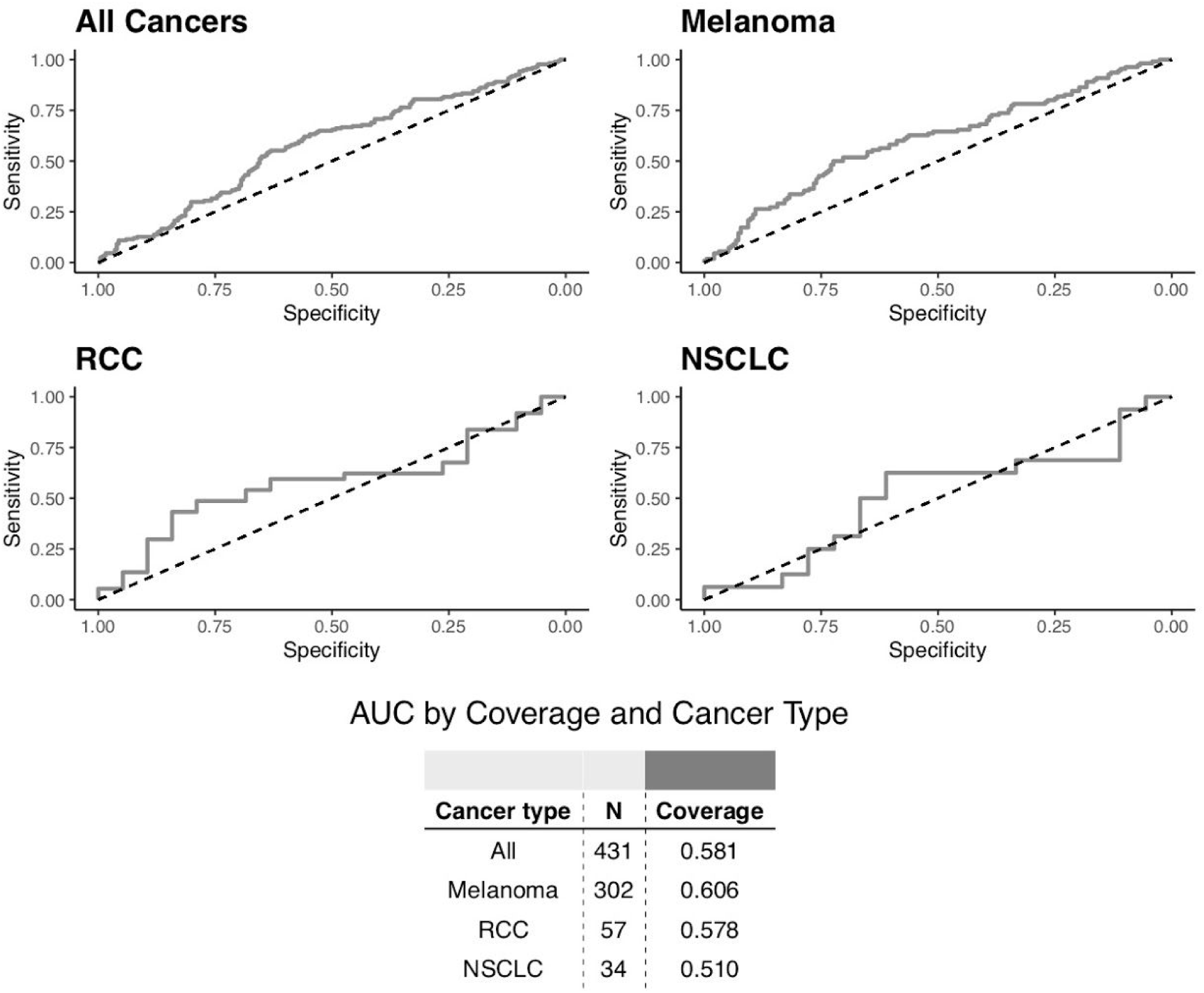
Receiver operating characteristic curves of predictive capacity of Mbp of genomic coverage. The upper panels depict the true positive rate (sensitivity, y-axis) and false positive rate (1-specificity, x-axis) for genomic coverage across all probability thresholds. The four panels represent models for four different cohorts based on different subsets of patients: All Cancers, which includes all patients, and Melanoma, RCC, and NSCLC, which include only melanoma, RCC, and NSCLC patients, respectively. The table in the lower panel reports the area-under-the-curve (AUC) for coverage (right column) applied to a different cancer cohort (rows). RCC=renal cell carcinoma, NSCLC=non-small cell lung cancer.

**Supplementary Figure 12:**
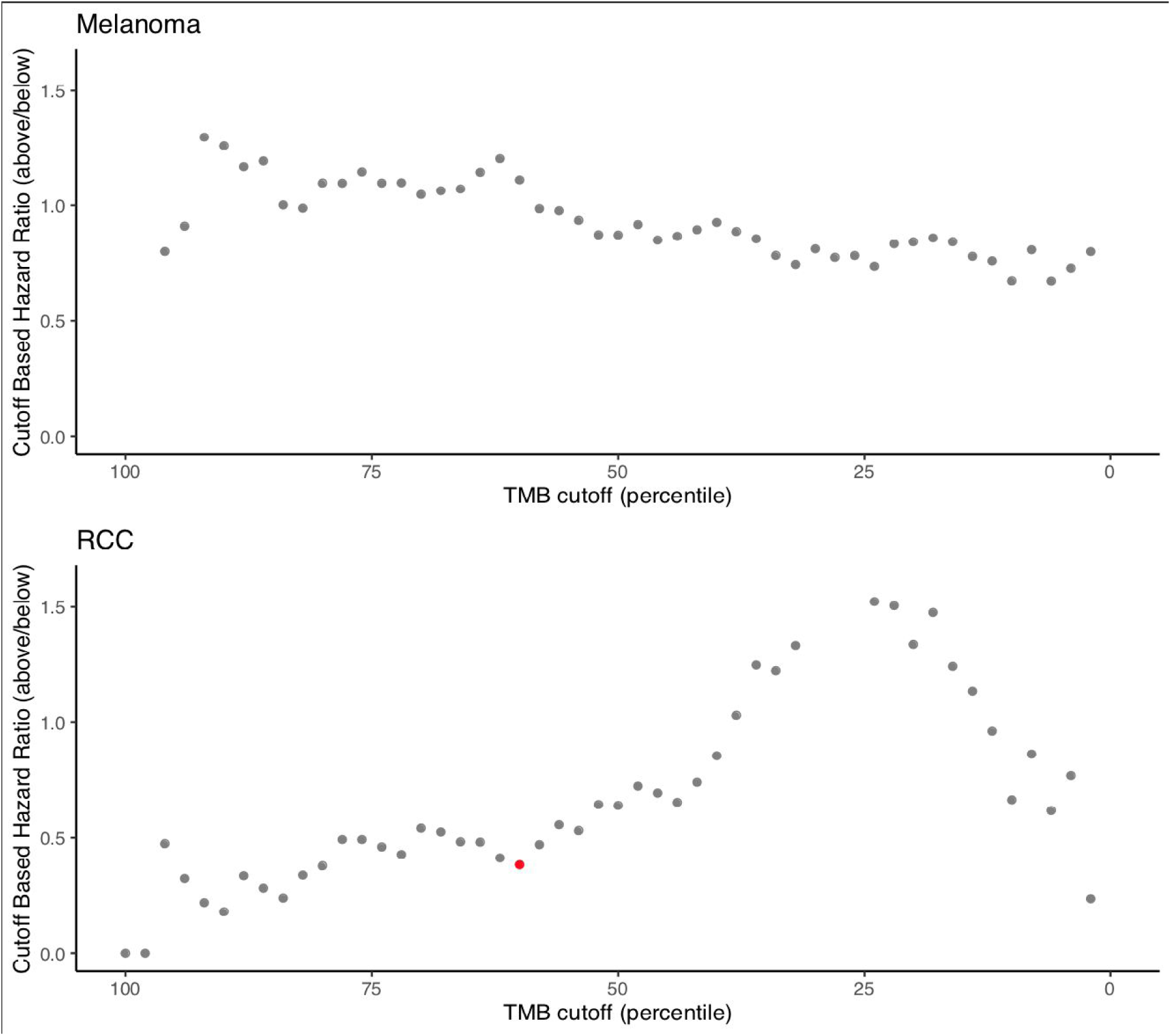
Variation in estimated hazard ratio based on TMB threshold selection. For melanoma and RCC separately, cox proportional hazard models were fit comparing patients above and below each TMB percentile cutoff at 2% intervals. The relative hazard ratio for those above the threshold compared to those below the threshold was plotted, with red representing models with corresponding unadjusted p-values < 0.05.

**Supplementary Figure 13:**
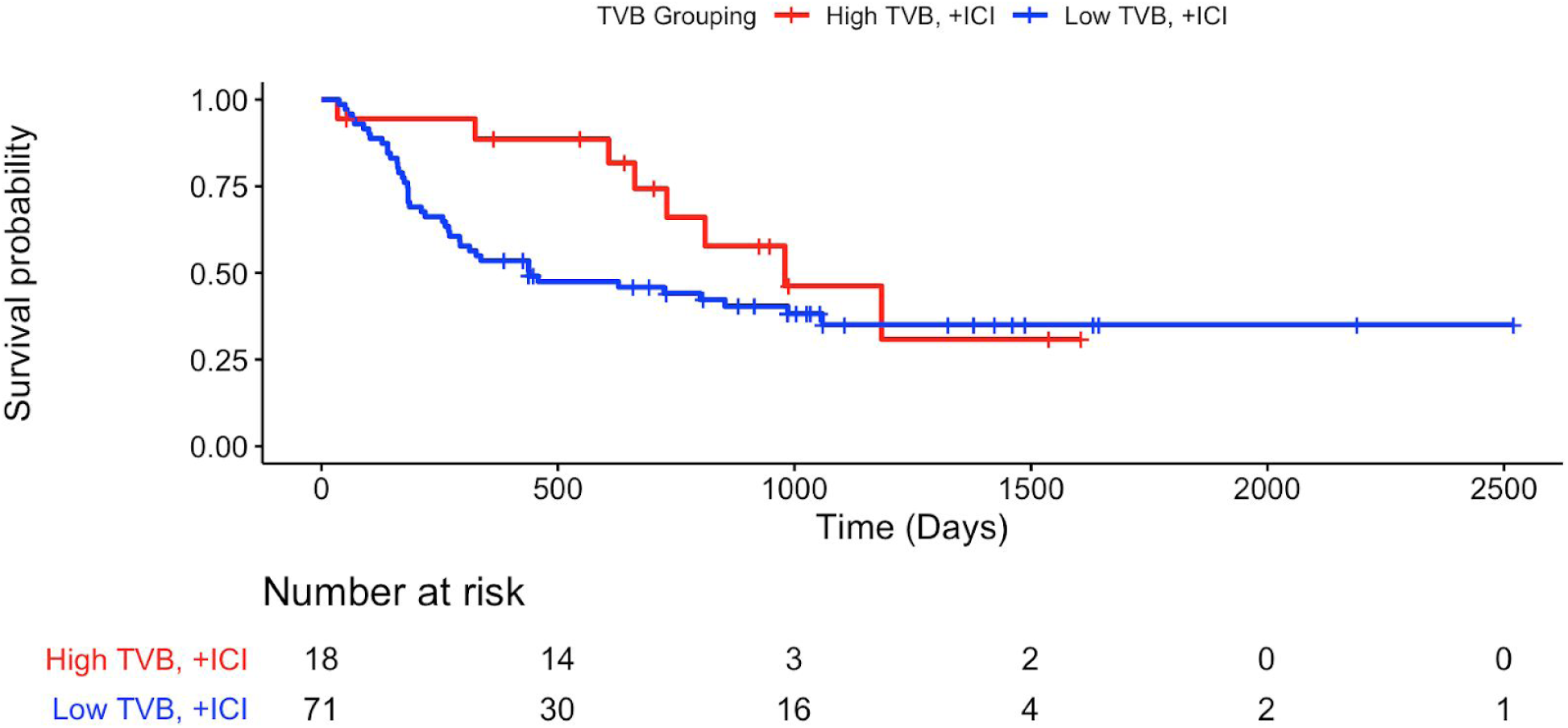
Overall survival among melanoma patients with high and low tumor variant burden (TVB). Kaplan-meier curves for the immunotherapy-treated patients with high TVB (≥80th percentile) and TVB burden (<80th percentile) are shown in red and blue, respectively. The underlying table corresponds to the number of patients at risk of death at each timepoint.

**Supplementary Figure 14:**
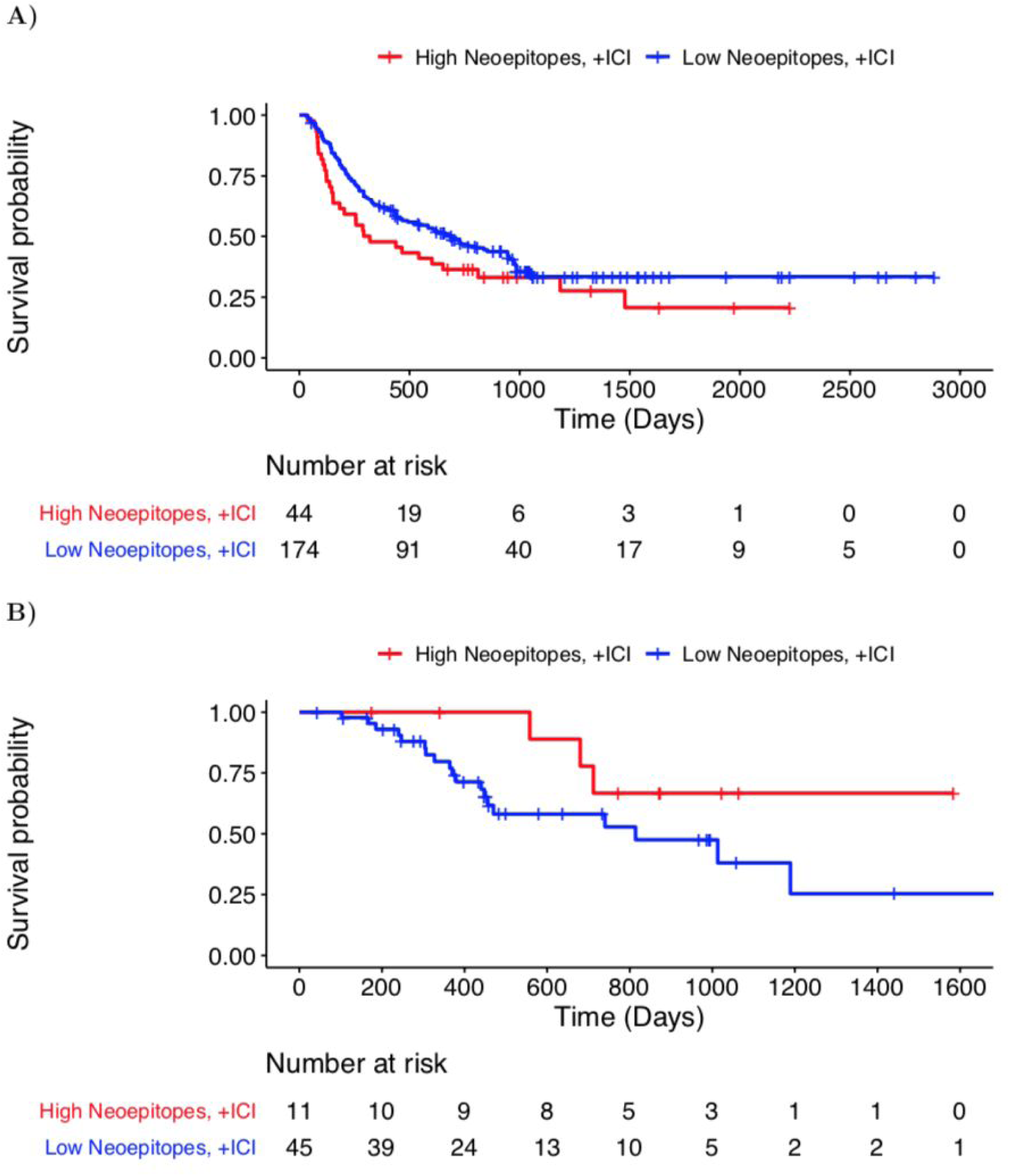
Overall survival among melanoma and renal cell carcinoma patients with high and low neoepitope burden. A) Overall survival among melanoma patients with high and low neoepitope burden. Kaplan-meier curves for the immunotherapy-treated patients with high neoepitope burden (≥80th percentile) and low neoepitope burden (<80th percentile) are shown in red and blue, respectively. The underlying table corresponds to the number of patients at risk for each timepoint. B) Overall survival among metastatic renal cell carcinoma patients with high and low neoepitope burdens. Kaplan-meier curves for the immunotherapy-treated patients with high neoepitope burden (≥80th percentile) and low neoepitope burden (<80th percentile) are shown in red and blue, respectively. The underlying table corresponds to the number of patients at risk for each timepoint.

**Supplementary Figure 15:**
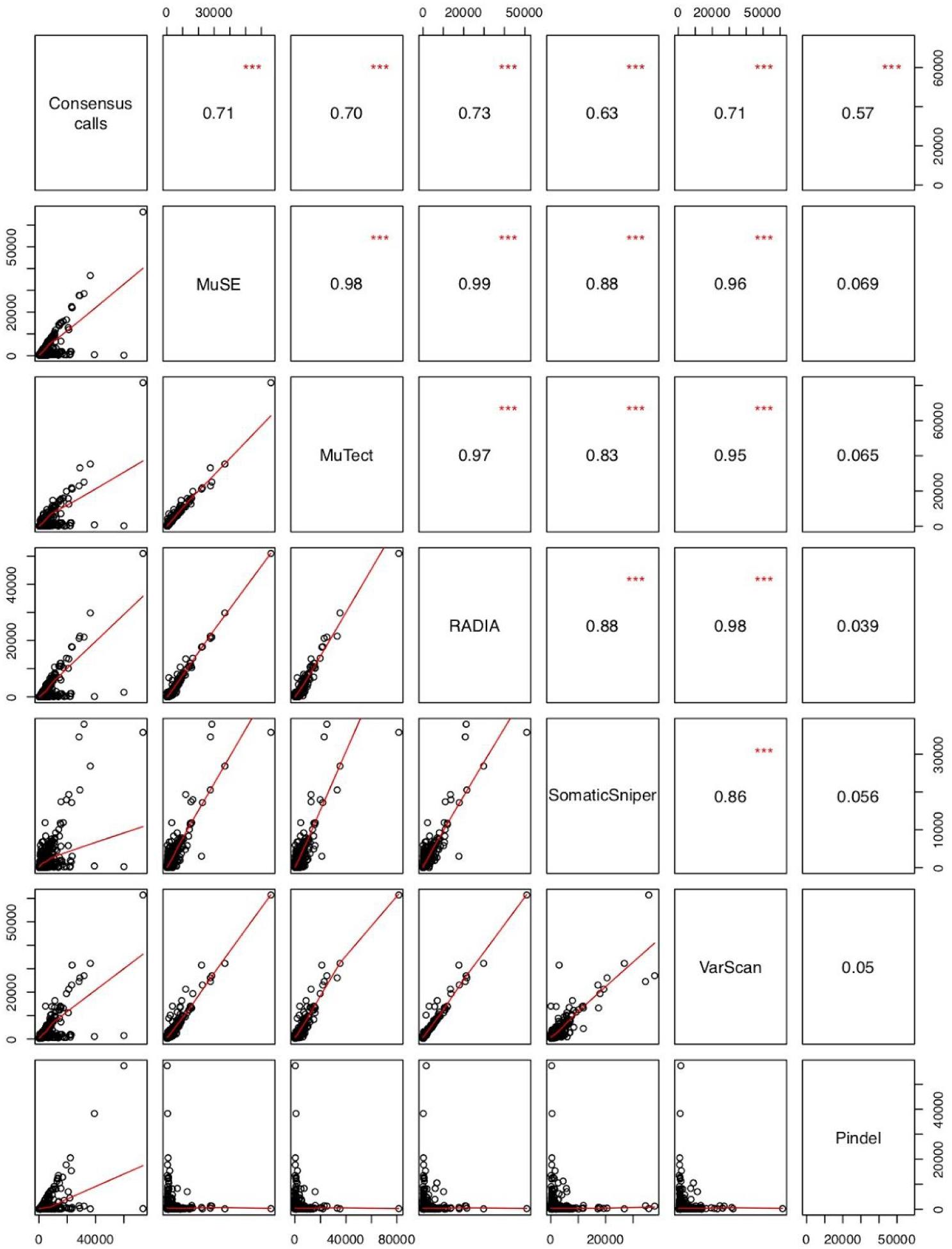
Pairwise differences in normalized total mutation burden as determined by 7 different computational approaches (see Methods). Each computational approach is identified along the diagonal panels, while the values in the upper panels denote the Pearson correlation coefficients between every pairwise combination of computational approaches (identified by corresponding row and column). The three red asterisks denote significant correlation at the p < 0.001 level. The scatterplots in the lower panels denote the TMB as calculated by each pairwise combination of computational approaches, with the x- and y-axes corresponding to the TMB calculated by the approach identified by the corresponding column and row, respectively; each open circle represents a single patient datapoint. Note that the red lines correspond to the best fit linear model.

**Supplementary Figure 16:**
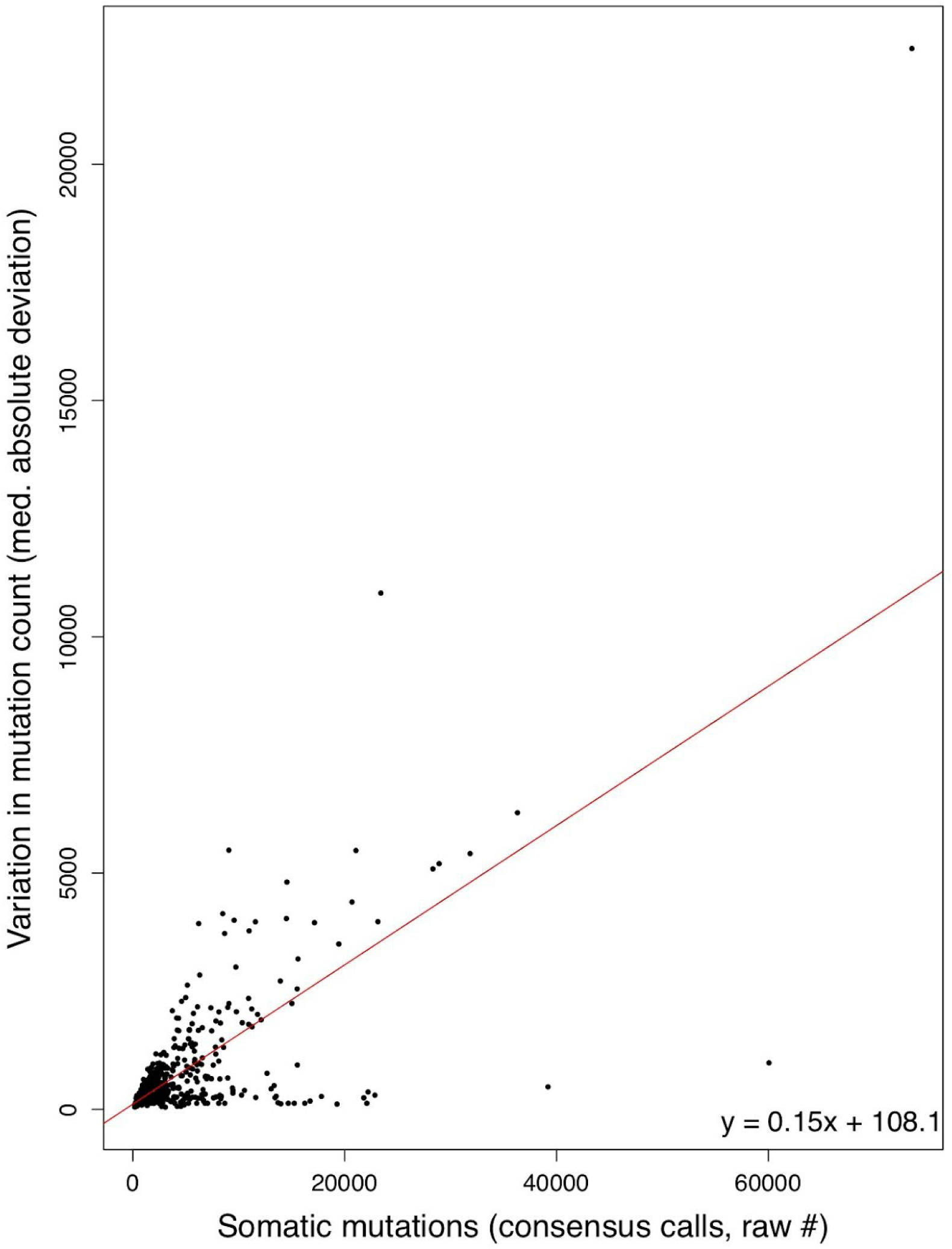
Variation in somatic mutation count increases with increased TMB from consensus variant calls. The median absolute deviation (MAD) in variant count across 6 variant calling tools used to determine consensus variant calls (y-axis, see Methods) increases with increasing TMB as determined by consensus calling (x-axis). The best fit line as determined by linear regression is shown in red, with its equation in the bottom right corner.

**Supplementary Figure 17:**
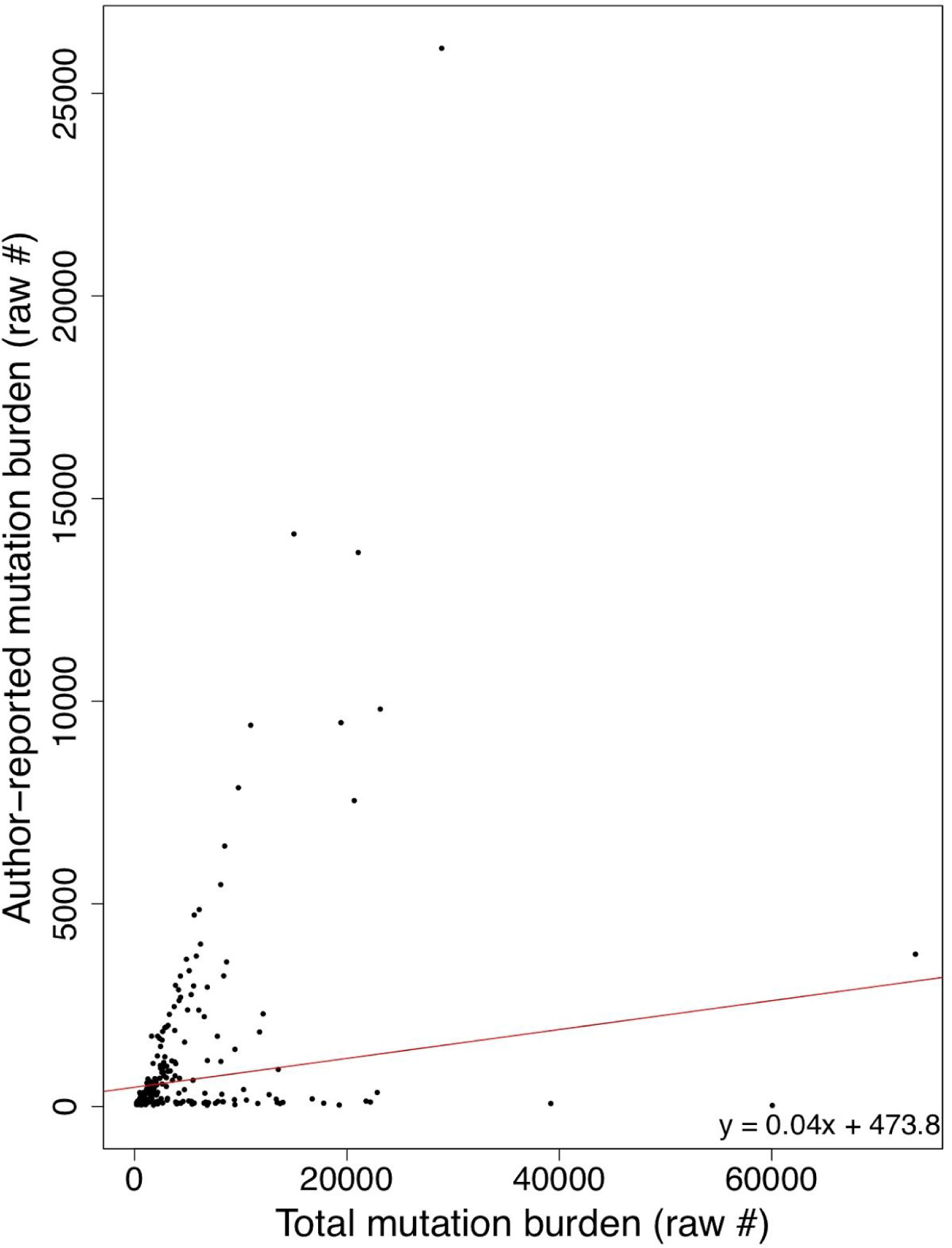
Author-reported total mutation burden correlates with consensus TMB. The total mutational burden as described by the authors of the original manuscripts from which the cohort derives (y-axis) correlates with our TMB derived from consensus variant calling (x-axis, Pearson product-moment correlation of 0.35, p = 1.99×10^−7^). The best fit line as determined by linear regression is shown in red, with its equation in the bottom right corner.

**Supplementary Figure 18:**
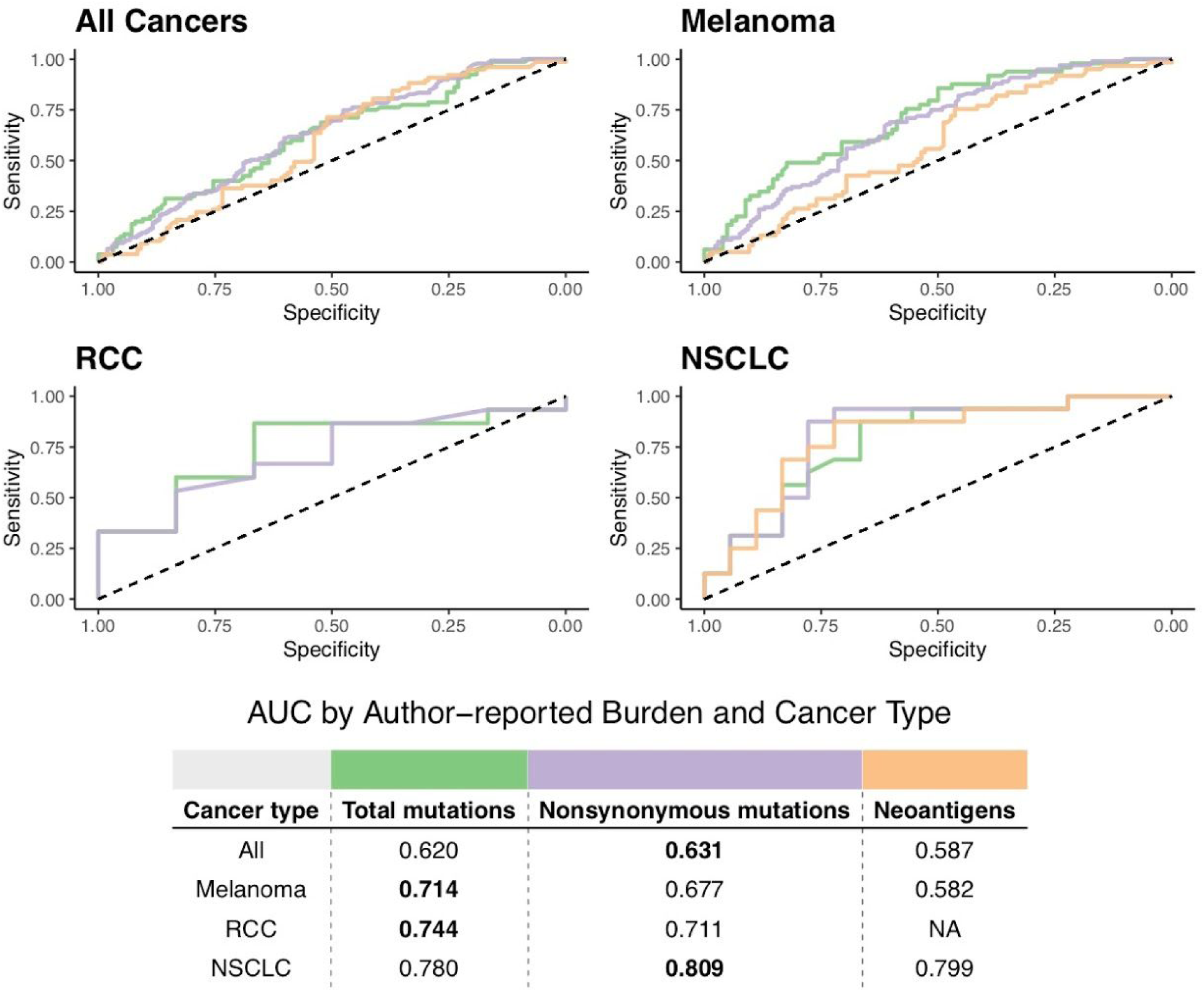
Receiver operating characteristic curves of predictive capacity of author-reported mutation and neoepitope burdens. The upper panels depict the true positive rate (sensitivity, y-axis) and false positive rate (1-specificity, x-axis) for each method across all probability thresholds. The four panels represent models for four different cohorts based on different subsets of patients: All Cancers, which includes all patients, and Melanoma, RCC, and NSCLC, which include only melanoma, RCC, and NSCLC patients, respectively. The table in the lower panel reports the area-under-the-curve (AUC) for each method (columns) applied to a different cancer cohort (rows), with colors above the methods indicating the color of the corresponding curve in the upper panels. Bold-faced values indicate the best value for each cancer cohort. RCC=renal cell carcinoma, NSCLC=non-small cell lung cancer.

**Supplementary Figure 19:**
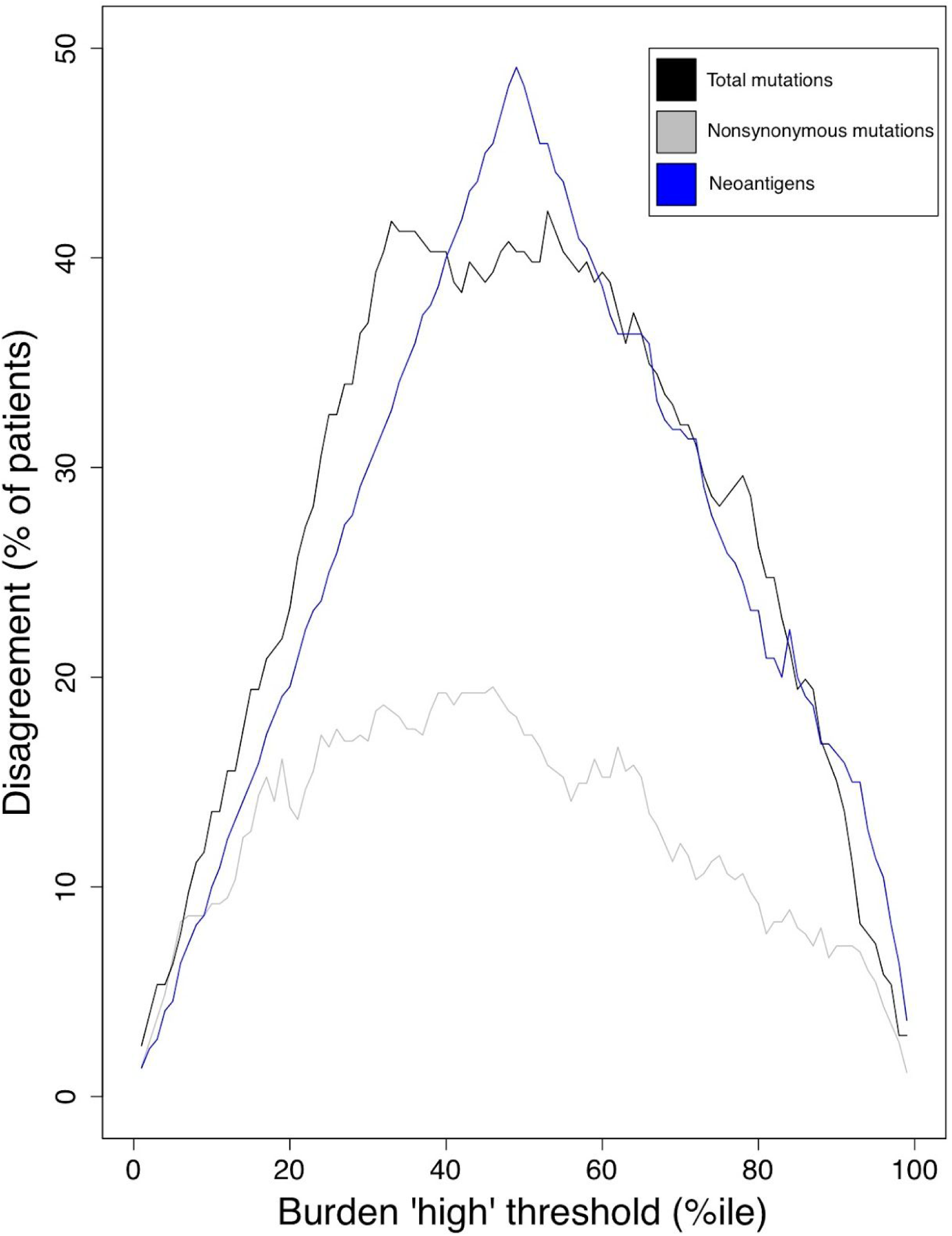
Cohort-level disagreement in classification of individual patients as TMB or neoepitope burden “high” v. “low”. TMB and neoepitope burdens were calculated using a standardized consensus approach (see Methods) and were compared with author-reported values from the original cohort source studies. The overall disagreement between classifications of consensus and author-reported data (y-axis) was calculated using different percentile thresholds (x-axis) to classify each individual as e.g. TMB “high” or “low”. This process was repeated for all mutations (black line), nonsynonymous mutations (gray line), and putative neoantigens (blue line).

**Supplementary Figure 20:**
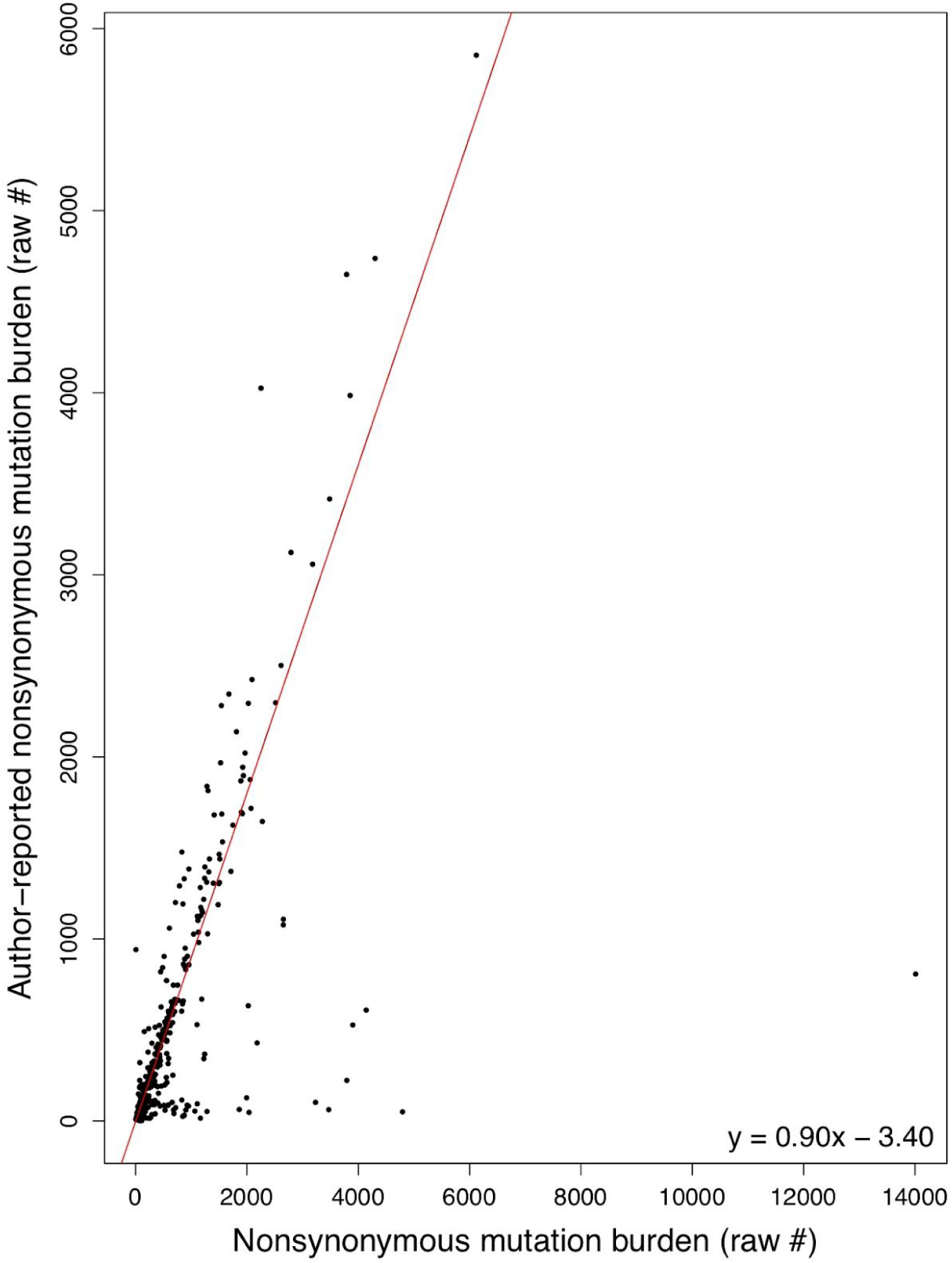
Author-reported nonsynonymous mutation burden correlates with nonsynoymous variants from consensus calling. The nonsynonymous mutational burden as described by the authors of the original manuscripts from which the cohort derives (y-axis) correlates with our consensus variant calling-derived nonsynonymous mutation burden (x-axis, Pearson product-moment correlation of 0.58, p < 2.2×10^−16^). The best fit line as determined by linear regression is shown in red, with its equation in the bottom right corner.

**Supplementary Figure 21:**
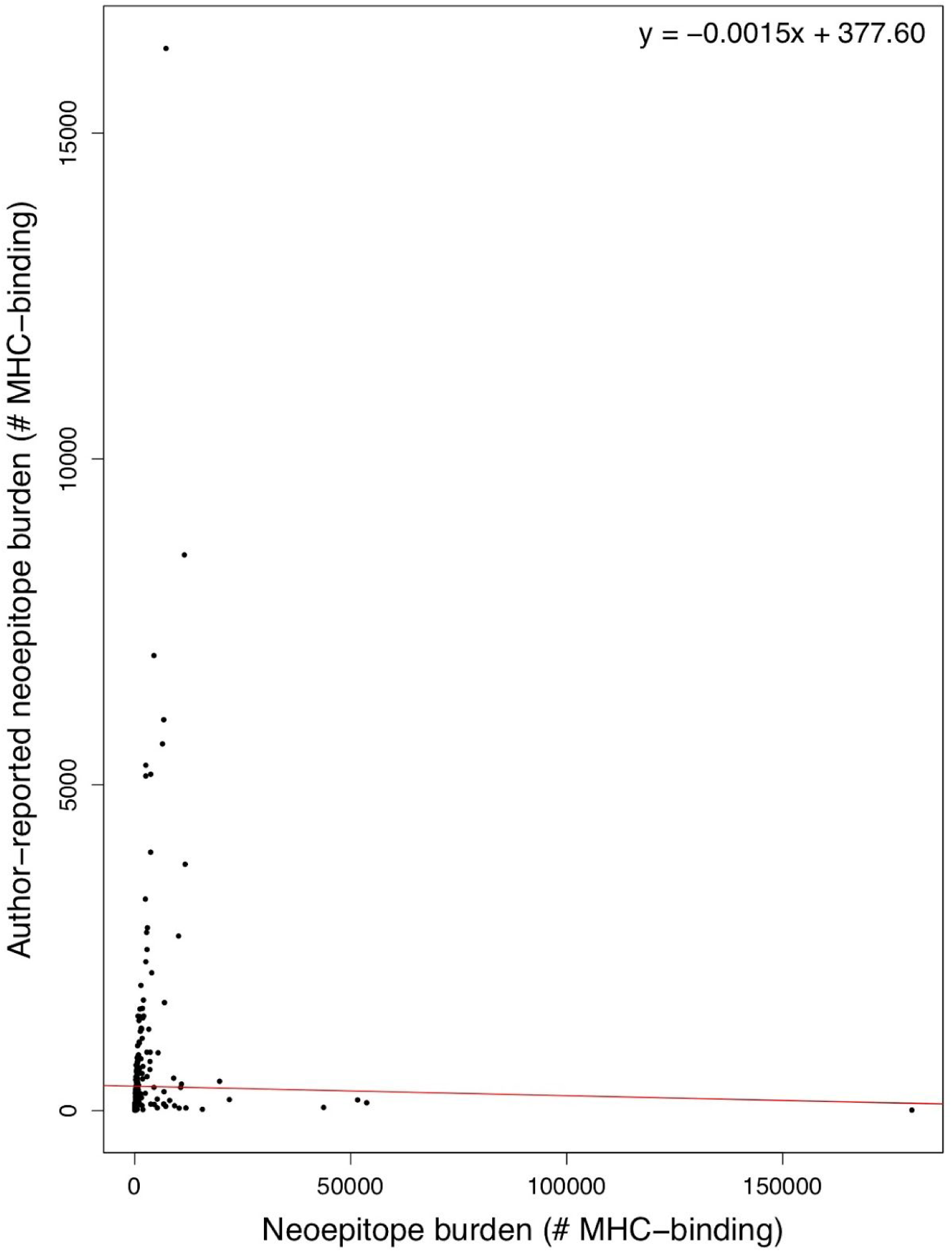
Author-reported neoepitope burden correlates with neoepitopes derived from variants from consensus calling. The neoepitope burden as described by the authors of the original manuscripts from which the cohort derives (y-axis) correlates with our consensus variant calling-derived neoepitope burden (x-axis, Pearson product-moment correlation of 0.026, p = 0.70). The best fit line as determined by linear regression is shown in red, with its equation in the bottom right corner.

**Supplementary Figure 22:**
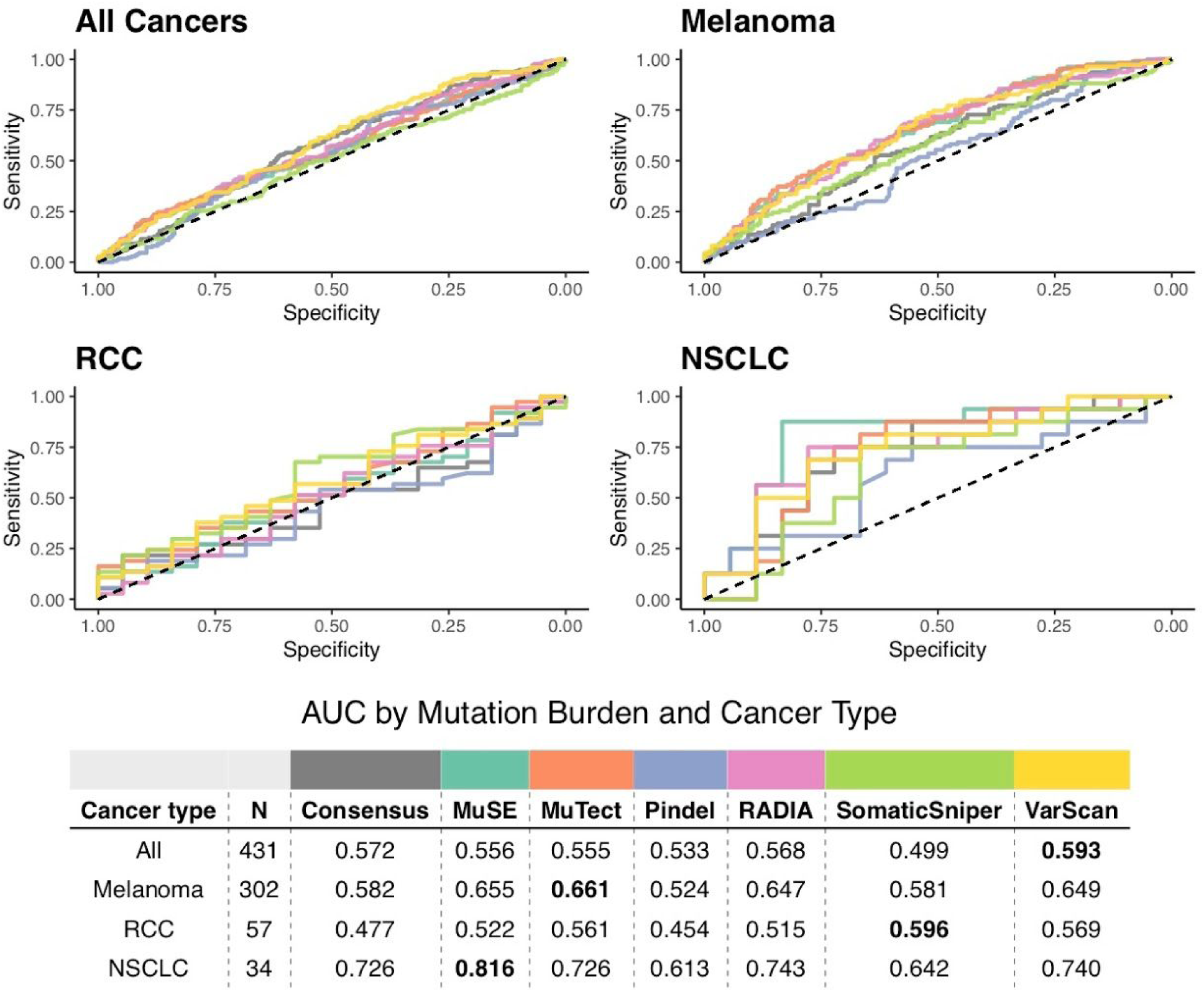
Receiver operating characteristic curves of predictive capacity of TMB from 7 different variant calling methods. The upper panels depict the true positive rate (sensitivity, y-axis) and false positive rate (1-specificity, x-axis) for each method across all probability thresholds. The four panels represent models for four different cohorts based on different subsets of patients: All Cancers, which includes all patients, and Melanoma, RCC, and NSCLC, which include only melanoma, RCC, and NSCLC patients, respectively. The table in the lower panel reports the area-under-the-curve (AUC) for each method (columns) applied to a different cancer cohort (rows), with colors above the methods indicating the color of the corresponding curve in the upper panels. TMB as determined by consensus calling (see Methods) is compared to the individual variant calling tools used in consensus calling. Bold-faced values indicate the best value for each cancer cohort. RCC=renal cell carcinoma, NSCLC=non-small cell lung cancer.

**Supplementary Figure 23:**
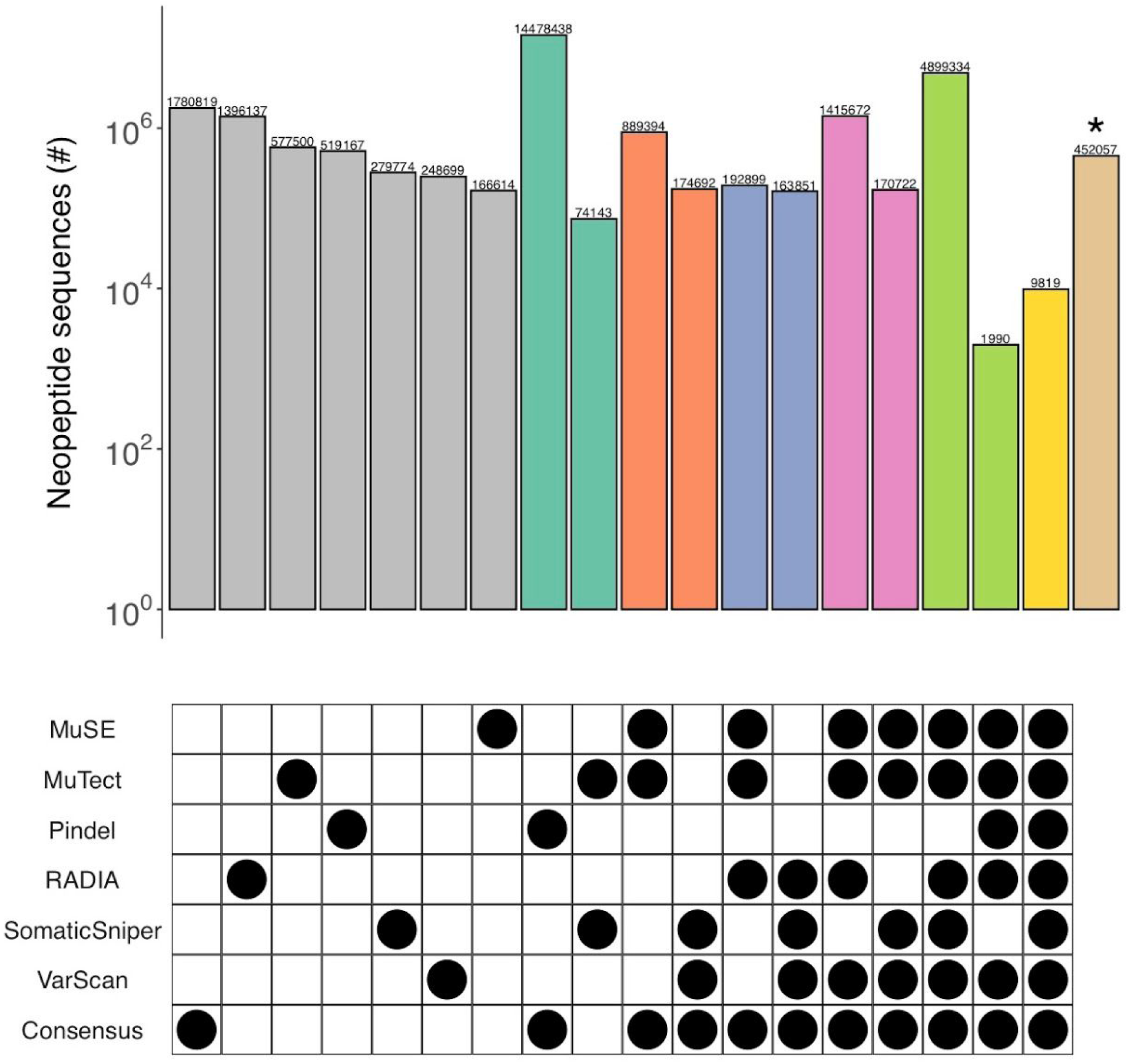
Detailed comparison of the complete set of neopeptide sequences predictions from MuSE, Mutect, Pindel, RADIA, SomaticSniper, VarScan, and consensus variant calling. Patterns of agreement or disagreement among groups of neopeptide sequences predicted from variants derived from different combinations of tools across all patients are shown along each column, and each row indicates the neopeptide predictions associated with variants from the indicated tool (e.g. the first column corresponds to neopeptides predicted only from Pindel variants). The number of neopeptides in each column (bar in upper pane) corresponds to the size of the subset predicted for variants from the indicated combination of tools (black circles in the bottom panel). Columns with gray bars represent neopeptides predicted from variants derived from only a single tool while columns with teal, orange, blue, pink, or green bars represent neopeptides predicted from variants derived from the most common two combinations of 2, 3, 4, 5, or 6 variant calling tools. The column with the yellow bar represents neopeptides predicted from variants deriving from all tools. The column with the brown bar (indicated by an asterisk) represents variants derived from less common combinations of 2-6 variant calling tools.

## REFERENCES

1. Van Allen EM, Miao D, Schilling B, Shukla SA, Blank C, Zimmer L, et al. Genomic correlates of response to CTLA-4 blockade in metastatic melanoma. Science. 2015;350(6257):207–11.

2. Samstein RM, Lee C-H, Shoushtari AN, Hellmann MD, Shen R, Janjigian YY, et al. Tumor mutational load predicts survival after immunotherapy across multiple cancer types. Nat Genet. 2019 Feb;51(2):202–6.

3. Rizvi NA, Hellmann MD, Snyder A, Kvistborg P, Makarov V, Havel JJ, et al. Cancer immunology. Mutational landscape determines sensitivity to PD-1 blockade in non-small cell lung cancer. Science. 2015 Apr 3;348(6230):124–8.

4. Carreno BM, Magrini V, Becker-Hapak M, Kaabinejadian S, Hundal J, Petti AA, et al. A dendritic cell vaccine increases the breadth and diversity of melanoma neoantigen-specific T cells. Science. 2015;348(6236):803–8.

5. Hugo W, Zaretsky JM, Sun L, Song C, Moreno BH, Hu-Lieskovan S, et al. Genomic and Transcriptomic Features of Response to Anti-PD-1 Therapy in Metastatic Melanoma. Cell. 2017 Jan 26;168(3):542.

6. Snyder A, Makarov V, Merghoub T, Yuan J, Zaretsky JM, Desrichard A, et al. Genetic basis for clinical response to CTLA-4 blockade in melanoma. N Engl J Med. 2014 Dec 4;371(23):2189–99.

7. Turajlic S, Litchfield K, Xu H, Rosenthal R, McGranahan N, Reading JL, et al. Insertion-and-deletion-derived tumour-specific neoantigens and the immunogenic phenotype: a pan-cancer analysis. Lancet Oncol. 2017 Aug;18(8):1009–21.

8. Smart AC, Margolis CA, Pimentel H, He MX, Miao D, Adeegbe D, et al. Intron retention is a source of neoepitopes in cancer. Nat Biotechnol. 2018;36(11):1056–8.

9. Le DT, Durham JN, Smith KN, Wang H, Bartlett BR, Aulakh LK, et al. Mismatch repair deficiency predicts response of solid tumors to PD-1 blockade. Science. 2017 Jul 28;357(6349):409–13.

10. Boyiadzis MM, Kirkwood JM, Marshall JL, Pritchard CC, Azad NS, Gulley JL. Significance and implications of FDA approval of pembrolizumab for biomarker-defined disease. J Immunother Cancer. 2018 May 14;6(1):35.

11. Meléndez B, Van Campenhout C, Rorive S, Remmelink M, Salmon I, D’Haene N. Methods of measurement for tumor mutational burden in tumor tissue. Transl Lung Cancer Res. 2018 Dec;7(6):661–7.

12. Zaretsky JM, Garcia-Diaz A, Shin DS, Escuin-Ordinas H, Hugo W, Hu-Lieskovan S, et al. Mutations Associated with Acquired Resistance to PD-1 Blockade in Melanoma. N Engl J Med. 2016 Sep 1;375(9):819–29.

13. Gao J, Shi LZ, Zhao H, Chen J, Xiong L, He Q, et al. Loss of IFN-γ Pathway Genes in Tumor Cells as a Mechanism of Resistance to Anti-CTLA-4 Therapy. Cell. 2016 Oct 6;167(2):397–404.e9.

14. Roh W, Chen P-L, Reuben A, Spencer CN, Prieto PA, Miller JP, et al. Integrated molecular analysis of tumor biopsies on sequential CTLA-4 and PD-1 blockade reveals markers of response and resistance. Sci Transl Med [Internet]. 2017 Mar 1;9(379). Available from: http://dx.doi.org/10.1126/scitranslmed.aah3560

15. Amaria RN, Reddy SM, Tawbi HA, Davies MA, Ross MI, Glitza IC, et al. Publisher Correction: Neoadjuvant immune checkpoint blockade in high-risk resectable melanoma. Nat Med. 2018 Dec;24(12):1942.

16. Eroglu Z, Zaretsky JM, Hu-Lieskovan S, Kim DW, Algazi A, Johnson DB, et al. High response rate to PD-1 blockade in desmoplastic melanomas. Nature. 2018 Jan 18;553(7688):347–50.

17. Graff JN, Alumkal JJ, Drake CG, Thomas GV, Redmond WL, Farhad M, et al. Early evidence of anti-PD-1 activity in enzalutamide-resistant prostate cancer. Oncotarget. 2016 Aug 16;7(33):52810–7.

18. Miao D, Margolis CA, Gao W, Voss MH, Li W, Martini DJ, et al. Genomic correlates of response to immune checkpoint therapies in clear cell renal cell carcinoma. Science. 2018 Feb 16;359(6377):801–6.

19. Hellmann MD, Nathanson T, Rizvi H, Creelan BC, Sanchez-Vega F, Ahuja A, et al. Genomic Features of Response to Combination Immunotherapy in Patients with Advanced Non-Small-Cell Lung Cancer. Cancer Cell. 2018 May 14;33(5):843–52.e4.

20. cancerit. cancerit/dockstore-cgpmap [Internet]. GitHub. [cited 2018 Sep 12]. Available from: https://github.com/cancerit/dockstore-cgpmap

21. Li H, Durbin R. Fast and accurate short read alignment with Burrows-Wheeler transform. Bioinformatics. 2009;25(14):1754–60.

22. gt. gt1/biobambam2 [Internet]. GitHub. [cited 2018 Sep 12]. Available from: https://github.com/gt1/biobambam2

23. Wood MA, Nguyen A, Struck A, Ellrott K, Nellore A, Thompson RF. neoepiscope improves neoepitope prediction with multi-variant phasing [Internet]. Bioinformatics. 2018. Available from: https://academic.oup.com/bioinformatics/advance-article/doi/10.1093/bioinformatics/btz653/5551338

24. Quinlan AR, Hall IM. BEDTools: a flexible suite of utilities for comparing genomic features. Bioinformatics. 2010 Mar 15;26(6):841–2.

25. Larson DE, Harris CC, Chen K, Koboldt DC, Abbott TE, Dooling DJ, et al. SomaticSniper: identification of somatic point mutations in whole genome sequencing data. Bioinformatics. 2012 Feb 1;28(3):311–7.

26. Koboldt DC, Zhang Q, Larson DE, Shen D, McLellan MD, Lin L, et al. VarScan 2: somatic mutation and copy number alteration discovery in cancer by exome sequencing. Genome Res. 2012 Mar;22(3):568–76.

27. Fan Y, Xi L, Hughes DST, Zhang J, Zhang J, Futreal PA, et al. MuSE: accounting for tumor heterogeneity using a sample-specific error model improves sensitivity and specificity in mutation calling from sequencing data. Genome Biol. 2016 Aug 24;17(1):178.

28. Cibulskis K, Lawrence MS, Carter SL, Sivachenko A, Jaffe D, Sougnez C, et al. Sensitive detection of somatic point mutations in impure and heterogeneous cancer samples. Nat Biotechnol. 2013 Mar;31(3):213–9.

29. Ye K, Schulz MH, Long Q, Apweiler R, Ning Z. Pindel: a pattern growth approach to detect break points of large deletions and medium sized insertions from paired-end short reads. Bioinformatics. 2009;25(21):2865–71.

30. Radenbaugh AJ, Ma S, Ewing A, Stuart JM, Collisson EA, Zhu J, et al. RADIA: RNA and DNA integrated analysis for somatic mutation detection. PLoS One. 2014 Nov 18;9(11):e111516.

31. Ellrott K, Bailey MH, Saksena G, Covington KR, Kandoth C, Stewart C, et al. Scalable Open Science Approach for Mutation Calling of Tumor Exomes Using Multiple Genomic Pipelines. Cell Syst. 2018 Mar 28;6(3):271–81.e7.

32. Tan A, Abecasis GR, Kang HM. Unified representation of genetic variants. Bioinformatics. 2015 Jul 1;31(13):2202–4.

33. McKenna A, Hanna M, Banks E, Sivachenko A, Cibulskis K, Kernytsky A, et al. The Genome Analysis Toolkit: a MapReduce framework for analyzing next-generation DNA sequencing data. Genome Res. 2010 Sep;20(9):1297–303.

34. Landrum MJ, Lee JM, Benson M, Brown GR, Chao C, Chitipiralla S, et al. ClinVar: improving access to variant interpretations and supporting evidence. Nucleic Acids Res. 2018 Jan 4;46(D1):D1062–7.

35. open-cravat [Internet]. Github; [cited 2019 May 30]. Available from: https://github.com/KarchinLab/open-cravat

36. Salipante SJ, Scroggins SM, Hampel HL, Turner EH, Pritchard CC. Microsatellite Instability Detection by Next Generation Sequencing [Internet]. Vol. 60, Clinical Chemistry. 2014. p. 1192–9. Available from: http://dx.doi.org/10.1373/clinchem.2014.223677

37. Dobin A, Davis CA, Schlesinger F, Drenkow J, Zaleski C, Jha S, et al. STAR: ultrafast universal RNA-seq aligner. Bioinformatics. 2013 Jan 1;29(1):15–21.

38. recount2: analysis-ready RNA-seq gene and exon counts datasets [Internet]. [cited 2019 May 15]. Available from: https://jhubiostatistics.shinyapps.io/recount/

39. GENCODE - Human Release 28 [Internet]. [cited 2019 May 15]. Available from: https://www.gencodegenes.org/human/release_28.html

40. Nellore A, Collado-Torres L, Jaffe AE, Alquicira-Hernández J, Wilks C, Pritt J, et al. Rail-RNA: scalable analysis of RNA-seq splicing and coverage. Bioinformatics. 2017 Dec 15;33(24):4033–40.

41. Bernstein MN, Doan A, Dewey CN. MetaSRA: normalized human sample-specific metadata for the Sequence Read Archive. Bioinformatics. 2017 Sep 15;33(18):2914–23.

42. MetaSRA [Internet]. [cited 2019 May 21]. Available from: http://metasra.biostat.wisc.edu/?and=CL:0000148%AC;=DOID:162&sampletype=cell%20line

43. MetaSRA [Internet]. [cited 2019 May 21]. Available from: http://metasra.biostat.wisc.edu/?and=CL:0000148%AC;=DOID:162&sampletype=primary%20cells

44. Wilks C, Gaddipati P, Nellore A, Langmead B. Snaptron: querying splicing patterns across tens of thousands of RNA-seq samples. Bioinformatics. 2018 Jan 1;34(1):114–6.

45. Snaptron User Guide — Snaptron 1.6 documentation [Internet]. [cited 2019 May 21]. Available from: http://snaptron.cs.jhu.edu/

46. kma [Internet]. Github; [cited 2019 May 13]. Available from: https://github.com/pachterlab/kma

47. Langmead B, Salzberg SL. Fast gapped-read alignment with Bowtie 2. Nat Methods. 2012 Mar 4;9(4):357–9.

48. Roberts A, Pachter L. Streaming fragment assignment for real-time analysis of sequencing experiments. Nat Methods. 2013 Jan;10(1):71–3.

49. NCI Primary Human Melanocyte QTL Study (ID 421623) - BioProject - NCBI [Internet]. [cited 2019 May 13]. Available from: https://www.ncbi.nlm.nih.gov/bioproject/PRJNA421623/

50. Szolek A, Schubert B, Mohr C, Sturm M, Feldhahn M, Kohlbacher O. OptiType: precision HLA typing from next-generation sequencing data. Bioinformatics. 2014 Dec 1;30(23):3310–6.

51. Boegel S, Löwer M, Schäfer M, Bukur T, de Graaf J, Boisguérin V, et al. HLA typing from RNA-Seq sequence reads. Genome Med. 2012 Dec 22;4(12):102.

52. Shao XM, Bhattacharya R, Huang J, Sivakumar IKA, Tokheim C, Zheng L, et al. High-throughput prediction of MHC Class I and Class II neoantigens with MHCnuggets [Internet]. Available from: http://dx.doi.org/10.1101/752469

53. Jurtz V, Paul S, Andreatta M, Marcatili P, Peters B, Nielsen M. NetMHCpan 4.0: Improved peptide-MHC class I interaction predictions integrating eluted ligand and peptide binding affinity data [Internet]. Available from: http://dx.doi.org/10.1101/149518

54. Stranzl T, Larsen MV, Lundegaard C, Nielsen M. NetCTLpan: pan-specific MHC class I pathway epitope predictions. Immunogenetics. 2010 Jun;62(6):357–68.

55. Altschul SF, Gish W, Miller W, Myers EW. Basic local alignment search tool. Journal of molecular [Internet]. 1990; Available from: https://www.sciencedirect.com/science/article/pii/S0022283605803602

56. Wood MA, Paralkar M, Paralkar MP, Nguyen A, Struck AJ, Ellrott K, et al. Population-level distribution and putative immunogenicity of cancer neoepitopes. BMC Cancer. 2018 Apr 13;18(1):414.

57. Tatlow PJ, Piccolo SR. A cloud-based workflow to quantify transcript-expression levels in public cancer compendia. Sci Rep. 2016 Dec 16;6:39259.

58. Lift Genome Annotations [Internet]. [cited 2019 May 13]. Available from: https://genome.ucsc.edu/cgi-bin/hgLiftOver

59. Broad GDAC Firehose [Internet]. [cited 2019 May 17]. Available from: http://gdac.broadinstitute.org/

60. Marty R, Kaabinejadian S, Rossell D, Slifker MJ, van de Haar J, Engin HB, et al. MHC-I Genotype Restricts the Oncogenic Mutational Landscape. Cell. 2017 Nov 30;171(6):1272–83.e15.

61. Ewing AD, Houlahan KE, Hu Y, Ellrott K, Caloian C, Yamaguchi TN, et al. Combining tumor genome simulation with crowdsourcing to benchmark somatic single-nucleotide-variant detection. Nat Methods. 2015 Jul;12(7):623–30.

62. Liu D, Schilling B, Liu D, Sucker A, Livingstone E, Jerby-Amon L, et al. Integrative molecular and clinical modeling of clinical outcomes to PD1 blockade in patients with metastatic melanoma. Nat Med. 2019 Dec;25(12):1916–27.

63. Chowell D, Morris LGT, Grigg CM, Weber JK, Samstein RM, Makarov V, et al. Patient HLA class I genotype influences cancer response to checkpoint blockade immunotherapy [Internet]. Vol. 359, Science. 2018. p. 582–7. Available from: http://dx.doi.org/10.1126/science.aao4572

64. Alspach E, Lussier DM, Miceli AP, Kizhvatov I, DuPage M, Luoma AM, et al. MHC-II neoantigens shape tumour immunity and response to immunotherapy. Nature. 2019 Oct;574(7780):696–701.

65. Chalmers ZR, Connelly CF, Fabrizio D, Gay L, Ali SM, Ennis R, et al. Analysis of 100,000 human cancer genomes reveals the landscape of tumor mutational burden. Genome Med. 2017 Apr 19;9(1):34.

66. Lyu G-Y, Yeh Y-H, Yeh Y-C, Wang Y-C. Mutation load estimation model as a predictor of the response to cancer immunotherapy. NPJ Genom Med. 2018 Apr 30;3:12.

67. Nguyen A, Garner C, Reddy SK, Sanborn JZ, Benz SC, Seery TE, et al. Three-fold overestimation of tumor mutation burden using 248 gene panel versus whole exome. J Clin Orthod. 2018 May 20;36(15_suppl):12117–12117.

68. Tumor Mutational Burden (TMB) [Internet]. Friends of Cancer Research. 2018 [cited 2019 May 29]. Available from: https://www.focr.org/tmb

69. Auslander N, Zhang G, Lee JS, Frederick DT, Miao B, Moll T, et al. Publisher Correction: Robust prediction of response to immune checkpoint blockade therapy in metastatic melanoma. Nat Med. 2018 Dec;24(12):1942.

70. Weiss GJ, Beck J, Braun DP, Bornemann-Kolatzki K, Barilla H, Cubello R, et al. Tumor Cell-Free DNA Copy Number Instability Predicts Therapeutic Response to Immunotherapy. Clin Cancer Res. 2017 Sep 1;23(17):5074–81.

71. Goodman AM, Kato S, Bazhenova L, Patel SP, Frampton GM, Miller V, et al. Tumor Mutational Burden as an Independent Predictor of Response to Immunotherapy in Diverse Cancers. Mol Cancer Ther. 2017 Nov;16(11):2598–608.

72. Anagnostou V, Niknafs N, Marrone K, Bruhm DC, White JR, Naidoo J, et al. Multimodal genomic features predict outcome of immune checkpoint blockade in non-small-cell lung cancer [Internet]. Vol. 1, Nature Cancer. 2020. p. 99–111. Available from: http://dx.doi.org/10.1038/s43018-019-0008-8

